# let-7 miRNA and *lin-46* mRNA are the two essential targets of the LIN28 RNA-binding protein in developmental timing

**DOI:** 10.64898/2026.01.12.699064

**Authors:** Jana Brunner, Anca Neagu, Dimos Gaidatzis, Lucas J. Morales Moya, Helge Großhans

## Abstract

The RNA-binding protein (RBP) LIN28 is a key regulator of temporal cell fates, promoting stem cell identity and suppressing differentiation. It regulates the processing of the let-7 miRNA but additionally binds thousands of mRNAs, suggesting a model of coordinated regulation across many targets. Yet, direct evidence for such a network function is scarce for both LIN28 and other broadly binding RBPs. Here, we show that in *C. elegans* larvae, loss of LIN28 leaves the levels of most LIN28-bound transcripts unchanged. The only exceptions are the *lin-46* mRNA and the let-7 miRNA. Comprehensive genetic analyses identify these two as the functionally essential LIN28 targets: *lin-28; lin-46; let-7* triple null mutant animals exhibit complete suppression of the *lin-28(0)* precocious heterochronic phenotypes and recapitulate the retarded *lin-46; let-7* double mutant phenotypes. Conversely, simultaneous overexpression of *lin-46* and *let-7* mimics LIN28 loss. Unexpectedly, LIN28 controls not only cell identity but also the tempo of development, using the same two targets. These findings establish a close functional link between the heterochronic pathway and the clock regulating developmental tempo and demonstrate that an RBP can achieve complex biological outcomes through a small set of targets.

## Introduction

Many animals undergo major morphological, physiological and/or behavioral changes as they become adults. Some of the genetic mechanisms that control this juvenile-to adult (J/A) transition, known as puberty in mammals, exhibit extensive evolutionary conservation (Faunes & Larrain, 2016; Moss & Romer-Seibert, 2014; Tena-Sempere, 2013). Thus, in *C. elegans*, heterochronic genes act as temporal patterning factors that determine stage-specific cell fates. Mutations in these genes cause either reiteration of earlier fates (retarded phenotypes) or skipping of earlier fates in favor of direct transition to a later fate (precocious phenotypes). A notable example is *lin-28*, which encodes an RNA-binding protein, and whose loss of function causes a precocious execution of the J/A transition (Ambros & Horvitz, 1984; Moss *et al*, 1997). Its mammalian orthologues *LIN28A* and *LIN28B* (collectively *LIN28* in the following) control puberty onset in mice and humans (Ong *et al*, 2009; Perry *et al*, 2009; Sulem *et al*, 2009; Zhu *et al*, 2010) (reviewed in (Cao *et al*, 2020)).

In both *C. elegans* and mammals, LIN28 functions, at least in part, by regulating biogenesis of the highly conserved miRNA let-7, preventing its precocious accumulation (Hagan *et al*, 2009; Heo *et al*, 2008; Lehrbach *et al*, 2009; Piskounova *et al*, 2011; Rybak *et al*, 2008; Van Wynsberghe *et al*, 2011; Viswanathan *et al*, 2008). At the cellular level, LIN28 and let-7 act antagonistically in controlling cell fates. While LIN28 promotes self-renewal, proliferation and “stemness”, let-7 drives differentiation. Together they form a regulatory module operating in diverse developmental, pathological and cell engineering contexts (Büssing *et al*, 2008; Tsialikas & Romer-Seibert, 2015).

Despite this important, phylogenetically conserved role of the LIN28–let-7 module, additional LIN28 targets must exist in *C. elegans*: In the absence of further targets, *let-7* would be epistatic to *lin-28*, i.e., *lin-28; let-7* double null mutant animals should exhibit the retarded phenotypes characteristic of *let-7* single mutant animals. Instead, these animals exhibit intermediate (or mutually suppressive) phenotypes. For instance, a cuticular structure known as adult alae form in wild-type animals at the fourth and final molt, M4. Loss of *lin-28* function causes precocious alae formation one or even two molts earlier, at M3 or M2, whereas *let-7* mutant animals fail to form alae even at M4. *lin-28; let-7* double mutant animals have partial alae at M4, but not before (Reinhart *et al*, 2000; Slack *et al*, 2000).

One additional LIN28 target is the *lin-46* mRNA, which LIN28 represses by binding to its 5’ untranslated region (5’UTR) (Ilbay *et al*, 2021). Like *let-7*, *lin-46* interacts genetically with *lin-28* and double mutations show mutual suppression (Pepper *et al*, 2004). However, many more targets are thought to exist: CLIP-seq revealed binding of LIN28 to more than 1,000 mRNAs, while yeast three-hybrid assays supported binding to precursors of the three let-7 sister miRNAs, miR-48, miR-84 and miR-241 (Stefani *et al*, 2015; Vadla *et al*, 2012).

Whether and to which extent these numerous interactions are functionally relevant remains mostly unknown. This is also true in mammalian cells where LIN28 was similarly reported to bind thousands of transcripts (Cho *et al*, 2012; Graf *et al*, 2013; Hafner *et al*, 2013; Wilbert *et al*, 2012). Although diverse molecular outcomes have been reported, including transcript stabilization, destabilization, translational repression or activation (Maklad *et al*, 2023), the observed effects are mostly modest and restricted to a fraction of the bound RNAs. Hence, it has been proposed that mRNA binding does not serve regulation of the mRNA, but sequesters LIN28 away from let-7, thereby permitting processing of the miRNA (Tan *et al*, 2019; Tan *et al*, 2021). Such a lack of individually important mRNA regulatory events could potentially explain why mammalian LIN28 RNA interactomes appear to differ greatly across different studies (Tsialikas & Romer-Seibert, 2015).

Here, we integrate RNA-binding and mRNA and miRNA expression data to identify LIN28-regulated RNAs in *C. elegans*. Surprisingly, among hundreds of bound mRNAs, only *lin-46* levels increase significantly in *lin-28* mutant animals, and among miRNAs, only let-7 is significantly upregulated. Consistent with these observations, genetic interaction studies reveal that LIN28’s somatic function in temporal patterning can be parsimoniously explained through only these two targets. Unexpectedly, loss of LIN28 also substantially slows overall development, dependent on derepression of *let-7* and *lin-46*. Hence, beyond acting as temporal patterning genes that organize developmental stages in a defined order, heterochronic genes may also control the tempo of development.

## Results

### LIN28 binding poorly predicts mRNA level changes

To identify functional targets of *C. elegans* LIN28, we performed RNA immunoprecipitation followed by sequencing (RIP-seq) by immunoprecipitating endogenously tagged LIN28. (We will refer to the relevant strain as *lin-28::large_tag* in the following to indicate that, in addition to the GFP epitope used in this experiment, it contains additional tags; Fig. 1A). Examining mid-L1 stage animals (6 h at 25°C after plating), we found 581 transcripts enriched for LIN28 binding (≥2-fold, Fig. 1B). Notably, this set overlapped significantly with targets identified by (Stefani *et al*., 2015) through HITS-CLIP (110/1015; 3.7-fold over expectation, *p* = 1.7 × 10⁻³⁴ hypergeometric test), despite differences in animal stage, genetic background, and experimental procedure.

**Figure 1.**
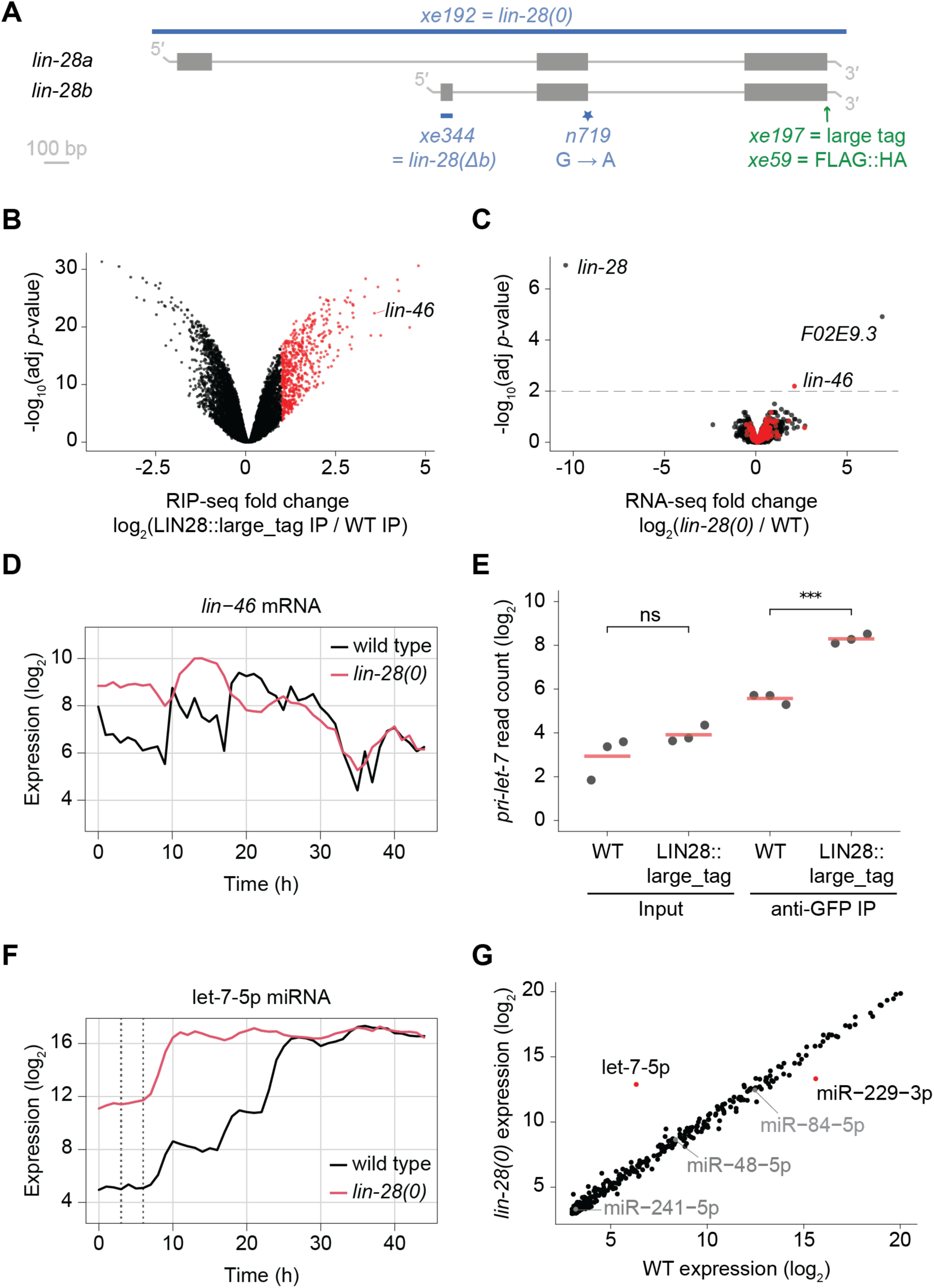
*let-7* and *lin-46* stand out among LIN28 candidate targets. **A**: Schematic representation of the *lin-28* genomic locus with two *lin-28* isoforms. Mutant alleles and endogenously tagged alleles used in this study are indicated. Allele numbers refer to modifications in otherwise wild-type backgrounds except the *lin-28b* isoform-specific exon deletion, *xe344*, which was introduced into an endogenously FLAG::HA-tagged background (*xe59*). The “large tag” *xe197* allele corresponds to a C-terminal insertion of linker::AID::3xFLAG::TEV::GFP::linker::HiBit sequence. **B**: Volcano plot showing transcript enrichment in LIN28::large_tag IP compared to untagged control IP (N2). Immunoprecipitation was performed with anti-GFP antibody on lysates from synchronized early L1-stage animals collected at 6 h of development at 25°C (n = 3 IP replicates). Transcripts significantly enriched (FDR ≤ 0.01, log_2_(FC) > 1) are indicated in red. **C**: Volcano plot showing differential gene expression in *lin-28(0)* animals relative to wild type. Log_2_ fold change in gene expression was quantified by RNA-seq of synchronized animals collected hourly from 3 to 6 h at 25°C in three biological replicates. The x-axis represents the average expression of each gene across the three biological replicates, with different time point samples pooled for each replicate. Three genes with significant expression changes (adjusted *p*-value < 0.01, threshold indicated by horizontal dashed line) are labeled by name. Genes whose transcripts are bound by LIN28 (from (**B**)) are indicated in red. *F02E9.3* expression was undetectable in both input and IP samples of LIN28::large_tag RIP-seq (shown in (**B**)). **D**: Expression levels of *lin-46* mRNA in wild-type and *lin-28(0)* animals as determined by mRNA-seq. Animals were grown at 25°C with hourly sampling. **E**: Plot showing log₂ read counts of *pri-let-7* in input and IP samples from LIN28::large_tag animals and untagged control animals (N2). Data are from the same LIN28::large_tag RIP-seq experiment as shown in (**B**). Red horizontal lines indicate the mean of three technical IP replicates. Significant differences were determined by an unpaired two-sided *t*-test (ns: not significant; \*\*\**p* < 0.001). **F**: Levels of mature let-7 miRNA in wild-type and *lin-28(0)* animals as determined by small RNA-seq. The two dashed lines indicate the window of time points pooled for differential gene expression analysis in (**G**). Samples are identical to those in (**D**). **G**: Scatterplot comparing miRNA expression in wild-type and *lin-28(0*) animals. miRNA expression levels were quantified by small RNA-seq of lysates from synchronized animals collected hourly from 3 to 6 h at 25°C. This corresponds to a subset of samples from the series shown in (**F**) but sequenced to greater depth and published previously in (Nahar *et al*, 2024). Samples were treated as pseudoreplicates for statistical testing, with the average miRNA expression across the four time points plotted. Two miRNAs with significant expression changes are colored in red (FDR < 0.01).

The role of LIN28 and its molecular mechanisms have previously been studied using *lin-28(n719)*, an allele isolated after chemical mutagenesis that carries a point mutation in the 5’ splice donor of the second exon and that is considered null (Ambros & Horvitz, 1984; Moss *et al*., 1997). To investigate the consequences of a complete loss of LIN28 in a genetically clean background, we introduced a ∼3.2 kb long deletion spanning the whole *lin-28* locus (*lin-28(xe192),* henceforth referred to as *lin-28(0)*, Fig. 1A). As we show below, *lin-28(0)* mutant animals recapitulate published *lin-28(n719)* mutant phenotypes (Ambros & Horvitz, 1984; Euling & Ambros, 1996), yet the two strains shared surprisingly few dysregulated genes (Fig. S1). Specifically, when compared to the wild-type N2 strain, many more genes were dysregulated in *lin-28(n719)* than in *lin-28(0)*, suggesting differences in strain background or additional mutations in the former.

**Figure S1.**
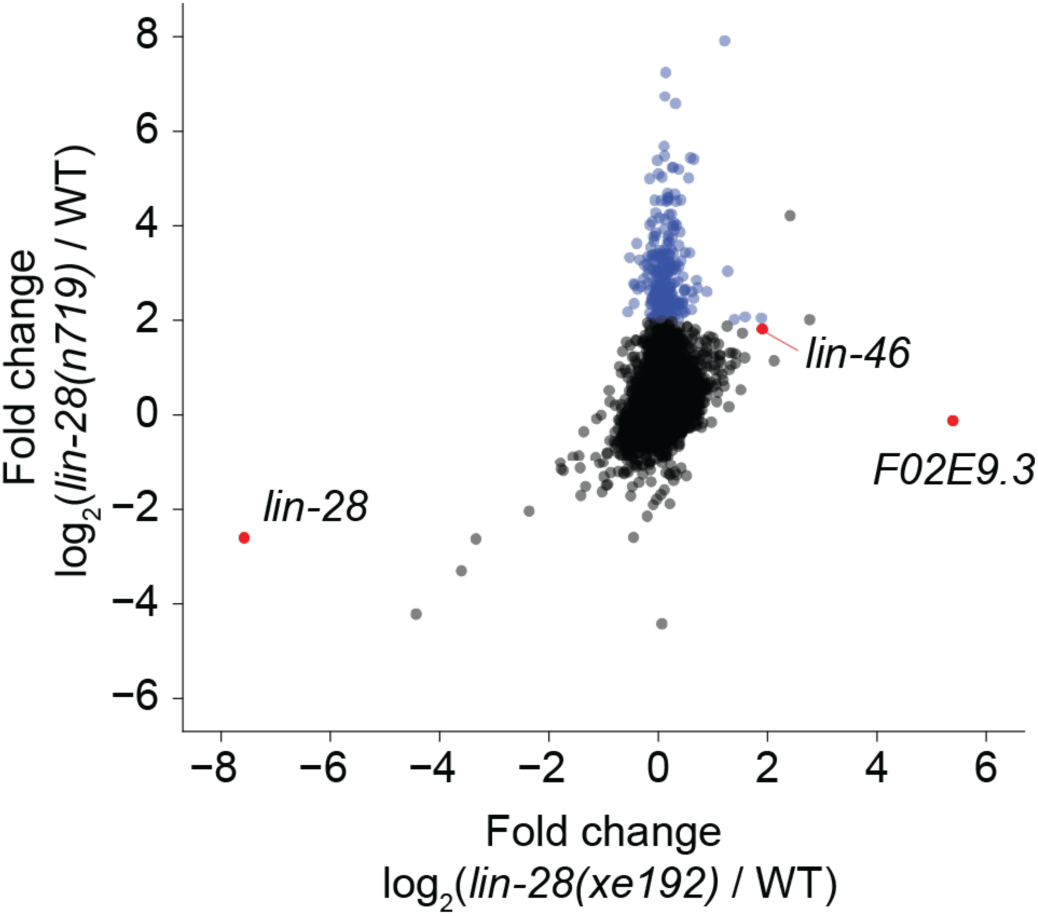
Gene expression profiles of *lin-28(n719)* and *lin-28(xe192)* differ substantially. Scatterplot depicting log_2_ fold changes in gene expression for each *lin-28* mutant relative to wild type. Expression was quantified by RNA-seq of synchronized animals collected hourly from 3 to 6 h at 25°C, with the average expression values across the four time points being shown. Independent biological replicates of the wild-type sample were used for the two comparisons. To highlight differences in global gene expression profiles, genes upregulated ≥4-fold in *lin-28(n719)* but ≤4-fold in *lin-28(xe192)* are labeled in blue. *lin-28*, *lin-46* and *F02E9.3* are labeled for reference. The changes in gene expression levels caused by the two *lin-28* alleles differ substantially, with only a handful of genes dysregulated in both backgrounds, including a previously validated LIN-28 target, lin-46 (Pepper *et al*., 2004). Moreover, while lin-28 mRNA is undetectable in *lin-28(xe192)* animals, its levels are only modestly reduced in *lin-28(n719)* relative to wild-type animals. Upon a closer inspection of the mapped sequencing reads, we noticed that in *lin-28(n719)* animals, an alternative splice site positioned 17 nucleotides upstream of the mutated one is used to splice the second and the third exon together, producing a lin-28 transcript harboring an in-frame stop codon. Strikingly, despite the substantial differences in their gene expression profiles, both *lin-28* mutations cause identical phenotypes, suggesting that the genes whose dysregulation causes the phenotypes are among the few affected by both lin-28 alleles.

To identify genes with robust changes in expression, we collected samples of several time points and in replicates for *lin-28(0)* and wild-type animals. We focused on the early first larval stage (eL1; 3 to 6 hours of development on food at 25°C), a time window when LIN28 levels are high (Moss *et al*., 1997; Seggerson *et al*, 2002) and potentially confounding rhythmic mRNA expression has not yet begun (Hendriks *et al*, 2014; Meeuse *et al*, 2020). For one of these experiments, sampling was extended to cover the entire course of larval development to allow assessment of mRNA trajectories over time. For those samples, we also sequenced small RNAs (see below).

Strikingly, besides *lin-28* mRNA itself, the levels of only two mRNAs changed significantly in *lin-28(0)* animals (Fig. 1C). The change in the first one, *F02E9.3*, is likely an artifact: It is not bound by LIN28 and its locus is immediately downstream of *lin-28*, suggesting that deletion of the *lin-28* coding sequence placed this gene under the control of the *lin-28* promoter. Consistent with this notion, *F02E9.3* levels were unaffected in *lin-28(n719)* animals (Fig. S1).

By contrast, the second transcript, *lin-46*, is not only bound by LIN28, but is also among the few transcripts whose levels were upregulated in both *lin-28(0)* and *lin-28(n719)* animals (Fig. S1). Moreover, regulation of *lin-46* by LIN28had previously been reported and was proposed to occur at the level of translation (Ilbay *et al*., 2021; Pepper *et al*., 2004). The long timecourse revealed that the levels of *lin-46* mRNA are dynamic in wild-type worms, with a notable increase during mid/late L1 (∼10 h) and again at ∼18 hours (∼mid-L2; Fig. 1D). In *lin-28(0)* animals, *lin-46* mRNA levels are upregulated during the first 18 hours of development (Fig. 1D), i.e., the time when LIN28 protein is abundant in wild-type animals (Moss *et al*., 1997; Seggerson *et al*., 2002). These observations suggest that LIN28 represses *lin-46*, at least in part, by destabilizing its transcript.

### let-7 is the only significantly upregulated miRNA in *lin-28* mutant animals

The best-characterized LIN28 target in *C. elegans* as elsewhere is the let-7 miRNA (Lehrbach *et al*., 2009; Nahar *et al*., 2024; Stefani *et al*., 2015; Van Wynsberghe *et al*., 2011), and our RIP-seq analysis confirmed binding of the primary let-7 transcript to LIN28 (Fig. 1E). (Note that Fig. 1B displays only mRNAs.) We had previously reported small RNA sequencing of the first 23 h of the long timecourse, which had revealed extensive upregulation of let-7 miRNA levels in *lin-28(0)* animals (Nahar *et al*., 2024). Interestingly, when we extended this analysis to the whole timecourse, we observed that the upregulation was specific to early larval development, whereas from ∼25 h onward, let-7 levels were identical in wild-type and *lin-28* mutant animals (Fig. 1F). This temporal pattern is consistent with the known restriction of LIN28’s activity to the early larval stages (Moss *et al*., 1997; Seggerson *et al*., 2002).

To examine whether LIN28 regulates the expression of other miRNAs, we determined differentially expressed miRNAs in the time window of 3 to 6 hours investigated also for mRNAs above. Using the four time points as pseudoreplicates for statistical testing, only let-7 and one other miRNA, miR-229-3p, changed significantly. In contrast to let-7, miR-229-3p levels were decreased, not increased, in *lin-28(0)* animals (Fig. 1G, Fig. S2). We expressly note that the let-7 sister miRNAs, miR-48, miR-84, and miR-241 were unaffected (Fig. 1G, Fig. S2).

**Figure S2.**
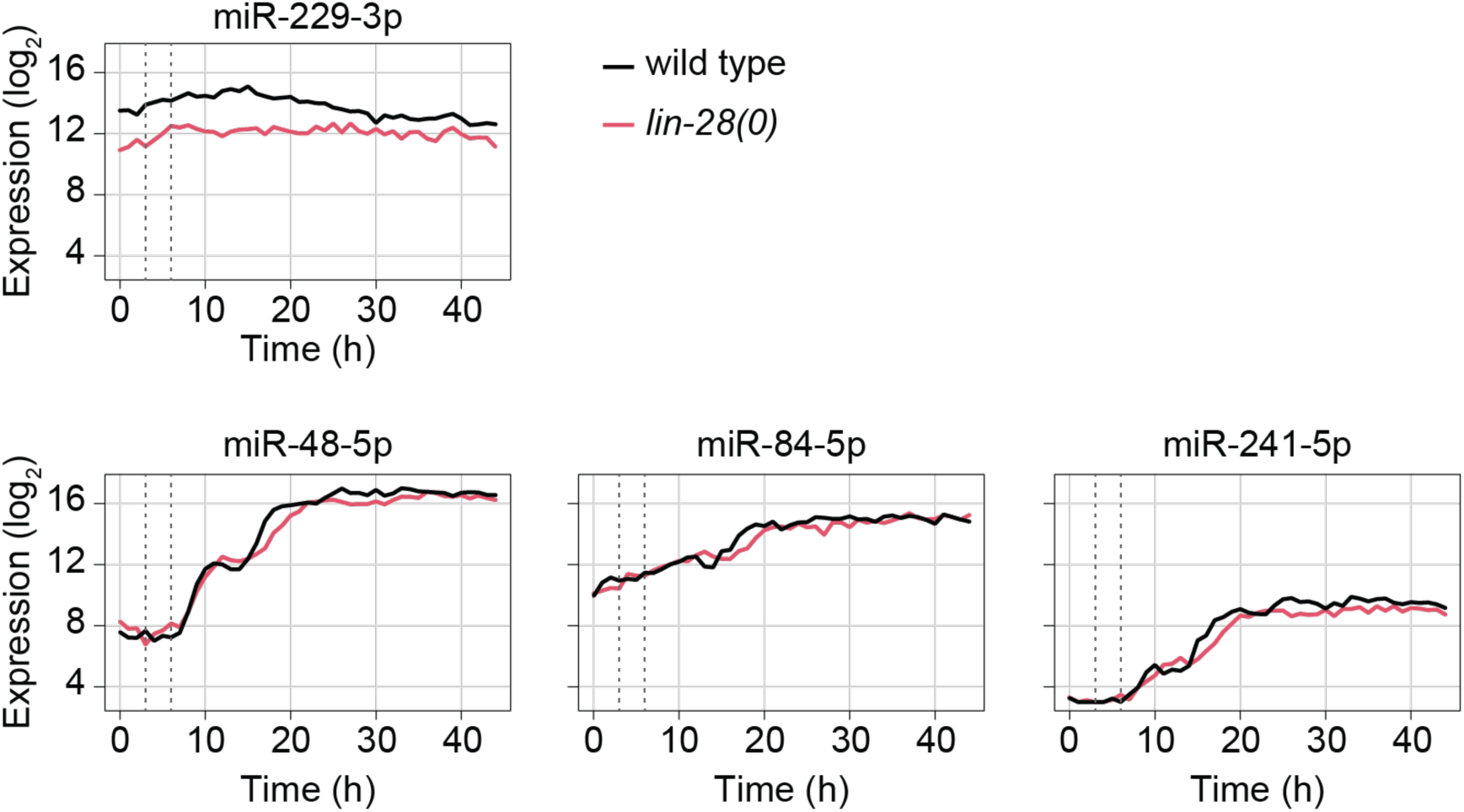
Developmental expression profiles of miRNAs in wild-type and *lin-28(0)* animals. Levels of mature miRNAs in wild-type and *lin-28(0)* animals as determined by small RNA-seq. The two dashed lines indicate the window of time points pooled for differential gene expression analysis in Fig. 1G.

Taken together, although LIN28 binds hundreds of transcripts, only a few of these binding events yield substantial changes in transcript levels. This suggests either that LIN28 acts preferentially through regulatory mechanisms other than RNA destabilization, or that most binding events be functionally dispensable. Hence, we sought to investigate to what extent regulation of only *let-7* and *lin-46* can account for the developmental functions of LIN28.

### *lin-46* but not *let-7* is required for the protruding vulva phenotype of *lin-28(0)* animals

Animals lacking LIN28 exhibit numerous heterochronic phenotypes, such as a protruding vulva due to precocious vulval precursor cell division, a precocious synthesis of the adult-specific skin structure called alae and a precocious cessation of the molting cycle after three instead of the wild-type four molts (Ambros & Horvitz, 1984; Euling & Ambros, 1996). We hypothesized that if *let-7* and *lin-46* were the only two developmentally relevant LIN28 targets, we should be able to explain these phenotypes by misregulation of just these two genes. In other words, mutating one or both target genes in the background of the *lin-28(0)* mutation should fully suppress the precocious *lin-28* mutant phenotypes. Moreover, the phenotypes resulting from their combined deletion should not be further modulated by loss of LIN28, their upstream repressor; i.e., *lin-28(0); lin-46(0); let-7(0)* triple mutant animals should recapitulate the retarded *lin-46(0); let-7(0)* double mutant phenotypes.

To test these two predictions, we used a previously published *lin-46(0)* allele, *lin-46(ma385)* (a ∼1.6 kb deletion spanning all *lin-46* exons; (Ilbay & Ambros, 2019)), and a newly generated *let-7(0)* allele, *let-7(xe150)*, which carries an ∼1 kb deletion of the promoter region (Fig. S3A) that prevents the synthesis of the mature let-7 miRNA (Fig. S3B).

**Figure S3.**
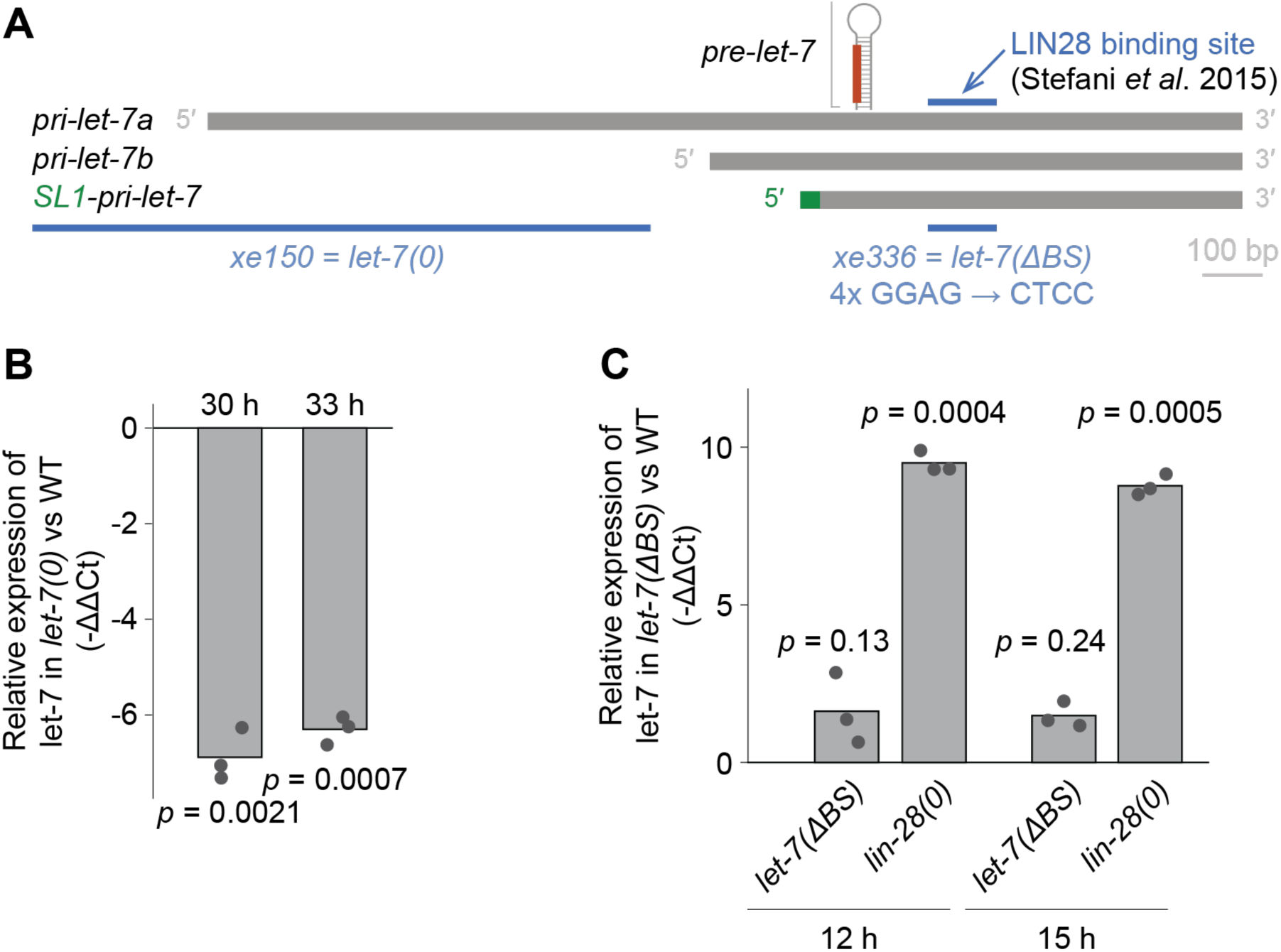
*let-7* alleles generated in this study. **A**: Schematic representation of the *let-7* genomic region depicting two isoforms of the let-7 primary transcript, *pri-let-7a* and *pri-let-7b*. Both isoforms undergo trans-splicing to spliced leader 1 RNA (SL1; shown in green), which is required for their further processing into mature let-7 (Bagga *et al*, 2005). The blue horizontal bar represents a putative LIN28 binding site (Stefani *et al*., 2015). The hairpin structure indicates the position of the *pre-let-7* hairpin within the primary transcripts, with the mature let-7 guide strand highlighted in red. Mutant alleles used in this study are indicated: *xe150* is a deletion, while *xe336* carries mutations converting the four GGAG motifs within the putative LIN28 binding site to CTCC. **B**: Mature let-7 levels were measured by TaqMan RT-qPCR in wild-type (N2) and *let-7(xe150)* animals at 30 h and 33 h of development at 25°C, when let-7 levels are at their peak in wild-type animals (Nahar *et al*., 2024). Expression levels at each time point were first normalized to the levels of small nucleolar RNA sn2841 (reference gene) and then to the wild-type sample. Bars indicate the mean of three biological replicates. *P*-values were determined by a paired two-sided *t*-test. **C**: Mature let-7 levels were measured by TaqMan RT-qPCR at 12 h and 15 h of development at 25°C, when LIN28 represses let-7 processing in wild-type animals. Expression levels at each time point were first normalized to the levels of small nucleolar RNA sn2841 (reference gene) and then to the wild-type sample. Bars indicate the mean of three biological replicates. *P*-values were determined by a paired two-sided *t*-test.

First, we examined vulval development. *lin-28(0)* animals exhibit a fully penetrant protruding vulva (Pvl) phenotype, which we find to be fully suppressed by the *lin-46(0)* allele (Fig. 2A), as reported earlier (Pepper *et al*., 2004). By contrast, this phenotype is not suppressed by a *let-7(0)* mutation (Fig. 2A). Instead, *lin-28(0)* suppresses a vulval phenotype associated with *let-7(0)*: *let-7(0)* mutant animals have a functionally deficient vulva through which they burst as young adults (Fig. 2B and (Reinhart *et al*., 2000)) – a phenotype that is absent from *lin-28(0); let-7(0)* double mutant animal (Fig. 2B). This finding can either be interpreted as *let-7* functioning upstream of *lin-28* or as evidence for *lin-28* regulating additional targets that act in parallel to *let-7*. Consistent with the latter scenario, we found that *lin-28(0); lin-46(0); let-7(0*) triple mutant animals phenocopied *let-7(0)* single mutant animals, exhibiting fully penetrant vulval bursting (Fig. 2B).

**Figure 2.**
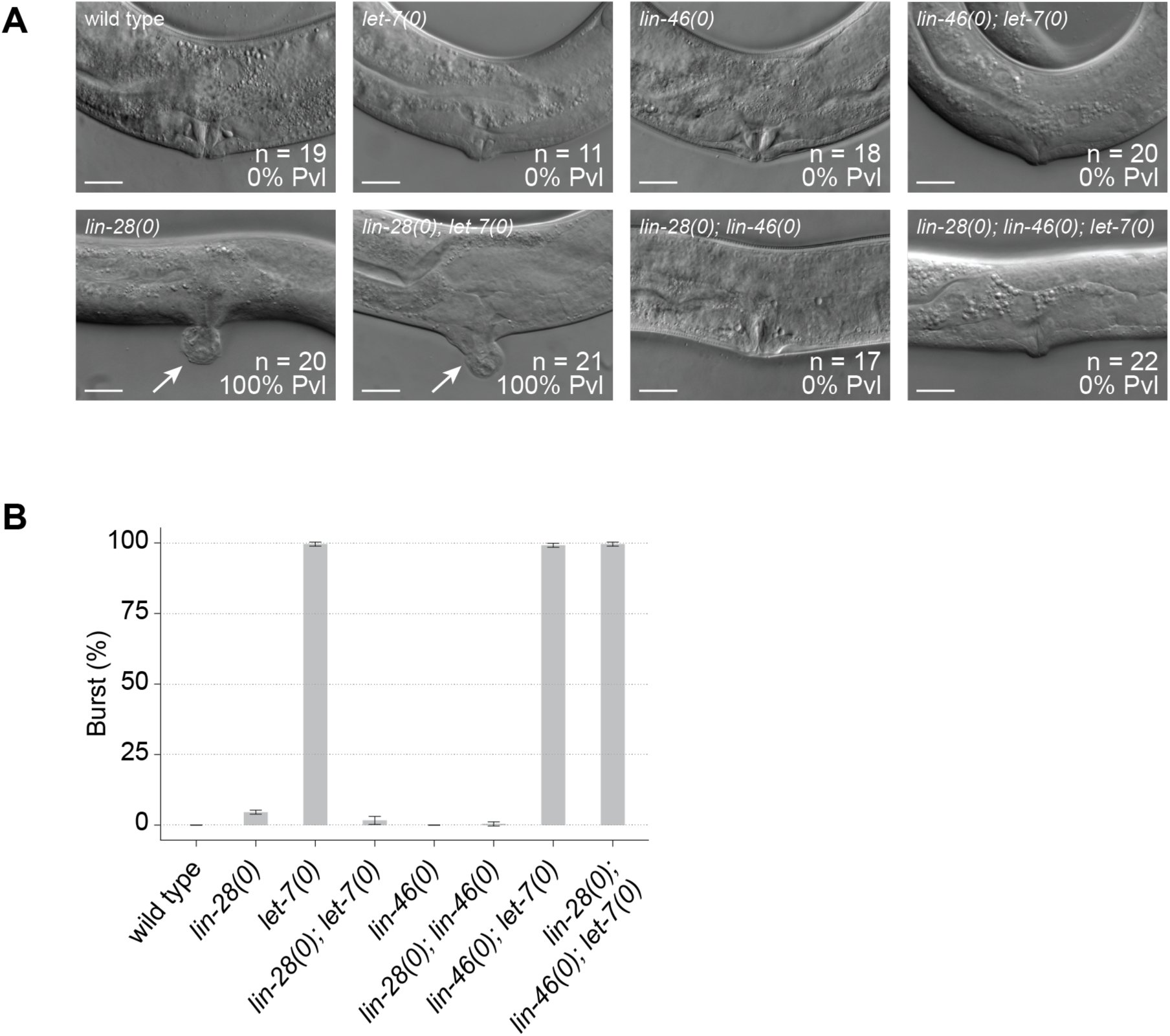
*lin-46* but not *let-7* is required for the *lin-28(0)* mutant protruding vulva (Pvl) phenotype. **A**: Example DIC images of vulvae in the indicated genetic backgrounds. The animals were grown at 20°C and examined shortly after exit from the final molt, i.e. the fourth molt in all cases except *lin-28(0)* animals, which undergo only three molts and were grown at 25°C. Arrow indicates the pathological vulva protrusion (Pvl). The percentage of animals exhibiting the Pvl phenotype is indicated. Scale bars: 10 µm. **B**: Quantification of vulva bursting of adult animals of indicated genotypes. Animals were grown at 25°C and examined at 48 h of development. N = 3 technical replicates, with n = 240 animals scored per genotype. Error bars represent standard deviation.

Taken together, our findings are consistent with *let-7* and *lin-46* acting in parallel pathways as targets of LIN28 with a partially redundant function in vulval development. Dysregulation of *lin-46* appears to chiefly account for the Pvl phenotype of *lin-28(0)* animals.

### Both *let-7* and *lin-46* are required for the precocious alae synthesis in *lin-28(0)* animals

Second, we evaluated the formation of alae, an adult-specific cuticular structure. In wild-type animals, alae are generated during the fourth molt. In contrast, *lin-28(0)* animals form alae precociously with 100% and 81% of animals having at least some alae already after the third molt when grown at 20°C and 25°C, respectively (Fig. 3A). In fact, and consistent with earlier work (Ambros & Horvitz, 1984), nearly half of the *lin-28(0)* animals exhibit weak partial alae as early as immediately after the second molt (Fig. 3A, B). The precocious alae synthesis is fully suppressed by loss of either *lin-46* or *let-7*, as neither *lin-28(0); lin-46(0)* nor *lin-28(0); let-7(0)* double mutant animals exhibit any alae after the third molt (Fig. 3A).

**Figure 3.**
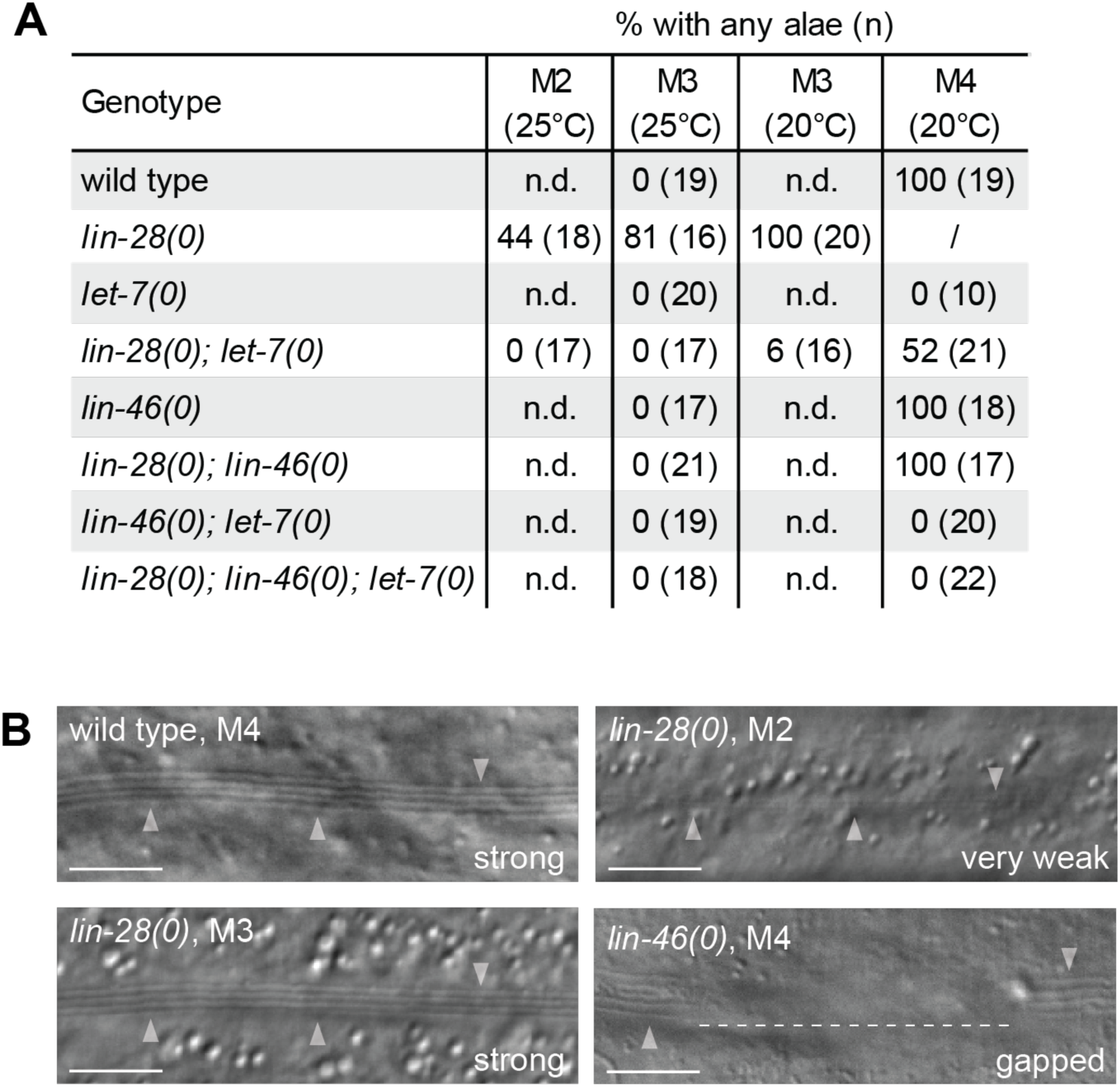
The precocious alae phenotype of *lin-28(0)* animals is suppressed by loss of either *let-7* or *lin-46*. **A**: Quantification of the presence of alae in animals of indicated genotypes. Molting animals were singled and examined shortly after they exited the respective molt as indicated by ecdysis. **B**: Example DIC images of alae in the indicated genetic backgrounds. The animals were grown at 20°C and examined shortly upon the exit from the indicated molt, with the exception of the “*lin-28(0)* M2” condition in which the animals were grown at 25°C. Arrowheads point to the alae. The dashed line indicates the alae gap. Scale bars: 10 µm.

Conversely, the *lin-28(0)* allele suppresses the alae phenotypes of the individual target mutations. Upon exit from the fourth molt, *lin-46(0)* animals have either complete (67%) or gapped (33%) alae, while all *lin-28(0); lin-46(0)* animals display complete, gap-free alae (Fig. 3A, B). At the same developmental stage, *let-7(0)* animals lack alae entirely, whereas around half of *lin-28(0); let-7(0)* animals exhibit weak partial alae (Fig. 3A). These findings again argue against a simple linear pathway where LIN28 controls alae formation through a single target. To determine whether regulation of *let-7* and *lin-46* is sufficient, we compared alae formation in *lin-46(0); let-7(0)* double and *lin-28(0); lin-46(0); let-7(0)* triple mutant animals. As in *let-7(0)* single mutant animals, neither strain formed any alae even after the fourth molt (Fig. 3A), supporting the conclusion that *let-7* and *lin-46* are likely the sole targets mediating LIN28’s role in the temporal control of alae synthesis. Moreover, unlike for the Pvl phenotype, dysregulation of both *lin-46* and *let-7* is required for precocious alae formation in *lin-28(0)* mutant animals.

### Both *let-7* and *lin-46* are required for the precocious exit from the molting cycle in *lin-28(0)* animals

Animals lacking LIN28 were previously reported to execute only three instead of the wild-type four larval molts (Ambros & Horvitz, 1984). To examine the contributions of *let-7* and *lin-46*, we employed a high-throughput assay allowing us to quantify the number of molts based on the alternating periods of feeding and lethargus in developing larvae (Meeuse *et al*., 2020; Olmedo *et al*, 2015). Briefly, animals expressing the luciferase transgene are provided with a medium containing luciferin, the luciferase substrate. As the animal grows and feeds, luciferin is ingested and, consequently, luminescence signal is emitted. When the animal enters a molt, it stops feeding and the signal decreases steeply, generating a characteristic luminescence intensity pattern (Fig. 4A).

**Figure 4.**
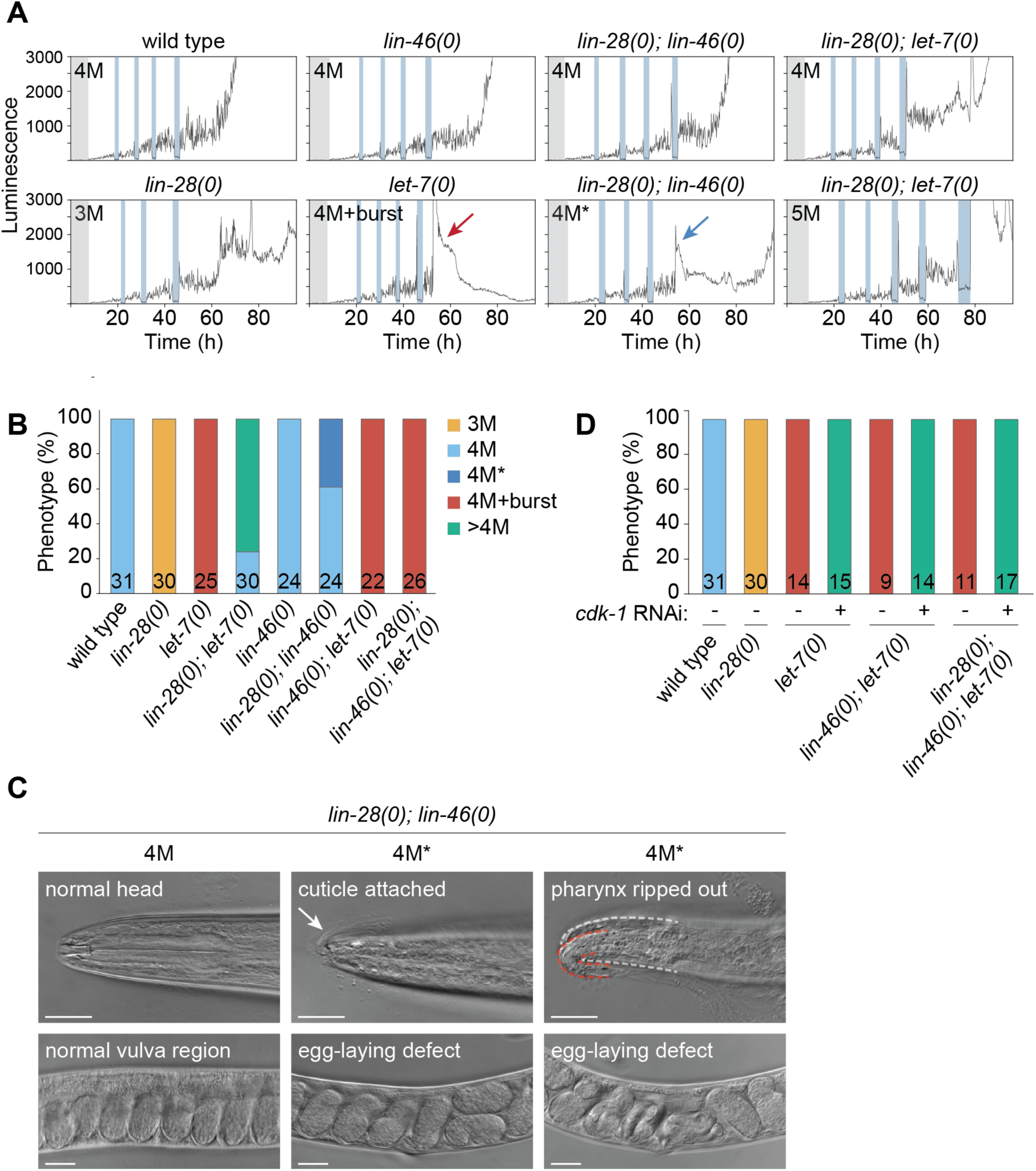
The precocious exit from the molting cycle of *lin-28(0)* animals is suppressed by both *let-7(0)* and *lin-46(0)* **A**: Examples of luminescence intensity traces. Vertical blue bars indicate molts. Red arrow points to the sudden signal increase indicative of vulva bursting. Blue arrow points to the sudden signal increase, the 4M* phenotype. **B**: Quantification of the number of molts in animals of the indicated genotypes. The number at the bottom of each bar indicates the number of examined animals (n). **C**: Example DIC images of the head (upper row) and the vulva (bottom row) regions *of lin-28(0); lin-46(0)* animals that underwent normal (left, 4M) or aberrant (middle and right, 4M*) fourth molt. The animals were extracted from the luciferase assay plate after 52 h, classified according to their luminescence intensity pattern and examined under the microscope. The 4M* animals exhibited a defect in pharyngeal cuticle shedding ranging from a part of the old cuticle remaining attached to the animal through the mouth plug (middle, white arrow) to a ripped-out pharynx (right, pharynx traced with a red dashed line, head traced with a white dashed line). All 4M* animals exhibited egg-laying defect as demonstrated by the retained embryos of advanced developmental stages. Scale bars: 10 μm. **D**: Quantification of the number of molts in animals of the indicated genotypes grown on either empty-vector containing (control) or *cdk-1* RNAi bacteria. Legend same as in (B). The wild type and *lin-28(0)* conditions are replotted from (B). The number at the bottom of each bar indicates the number of examined animals (n).

The analysis revealed the expected four molts in wild-type and three in *lin-28(0)* animals (Fig. 4A, B). The precocious exit from the molting cycle observed upon LIN28 loss requires the activities of both *let-7* and *lin-46*, as both *lin-28(0); let-7(0)* and *lin-28(0); lin-46(0)* double mutant animals undergo more than three molts (Fig. 4A, B). However, in the case of *lin-28(0); lin-46(0)* double mutants, suppression was only partial: nearly half of the animals exhibit a defect during the fourth molt (henceforth referred to as the 4M* phenotype) (Fig. 4B): After completing three properly executed molts, a steep increase in the luminescence intensity occurred at a time where wild-type animals would exhibit the signal drop typical of feeding cessation (Fig. 4A). This unusual signal reflects a severe defect in shedding of the pharyngeal cuticle, with the anterior part of the old cuticle remaining attached via the mouth plug (Fig. 4C, middle). In extreme cases, the pharynx appears to be partially ripped out, likely due to mechanical forces applied during failed cuticle shedding (Fig. 4C, right). Additionally, late-stage embryos accumulate in the affected animals, indicating an egg-laying defect (Fig. 4C), possibly a consequence of embryo retention due to starvation (Chen & Caswell-Chen, 2004). We note that the 4M* phenotype is specific to liquid culture as *lin-28(0); lin-46(0)* animals appear wild-type when grown on agar plates.

*lin-46(0)* mutant animals undergo the wild-type number of four molts. By contrast, if *let-7(0)* animals are prevented from vulval bursting, they execute additional molts. Thus, 100% of *let-7(0)* animals depleted of CDK-1 to prevent vulva formation (Rausch *et al*, 2015) undergo extra molts (Fig. 4D). The *lin-28(0)* mutation not only fully suppressed vulval bursting, but also partially suppressed the extra molt phenotype with 24% of *lin-28(0); let-7(0)* double mutant animals undergoing the normal number of molts, four (Fig. 4A, B). This suppression depended on *lin-46*: Like *let-7(0)* single and *lin-46(0); let-7(0)* double mutant animals, all *lin-28(0); lin-46(0); let-7(0)* triple mutant animals executed supernumerary molts when grown on *cdk-1* RNAi (Fig. 4C). We conclude that *let-7* and *lin-46* are the essential targets of LIN28 for controlling molt number. As for the alae phenotype, precocious termination of the molting cycle in *lin-28(0)* mutant animals appears to require dysregulation of both *lin-46* and *let-7*.

### Mutation of the putative LIN28 binding site in the primary let-7 transcript does not alter let-7 levels

With the suppression analysis we tested whether the two LIN28 targets, *let-7* and *lin-46*, are necessary for *lin-28* mutant phenotypes. Next, we wanted to test whether the precocious expression of these targets is sufficient to induce those phenotypes. Thus, we aimed to use genome editing to uncouple the two targets from LIN28-imposed repression.

In humans, LIN28 recognizes a GGAG motif located in the apical part of the *let-7* hairpin (Loughlin *et al*, 2011; Nam *et al*, 2011; Ustianenko *et al*, 2018). However, the *let-7* hairpin in *C. elegans* lacks this motif, and instead, LIN28 has been proposed to bind a downstream region outside of the actual hairpin ((Stefani *et al*., 2015); Fig. S3A). This region contains four GGAG motifs, which have been implicated in LIN28 binding *in vitro* (Stefani *et al*., 2015). To uncouple let-7 from LIN28-mediated repression *in vivo*, we mutated all four motifs in the endogenous *let-7* locus, changing them from GGAG to CTCC (*let-7(xe336)*, henceforth referred to as *let-7(ΔBS)* for mutation of the putative <u>B</u>inding <u>S</u>ites) (Fig. S3A). However, this mutation did not result in appreciable *let-7* derepression, as mature let-7 levels remained largely unaffected (Fig. S3C), suggesting that the four GGAG motifs in the putative LIN28 binding site are not required for let-7 repression in vivo. (We have not determined whether LIN28 binding was perturbed by these mutations.)

### Mutation of the putative LIN28 binding site in *lin-46* 5’UTR results in weakly precocious LIN-46 accumulation

The *lin-46* transcript is bound by LIN28 (Fig. 1B and (Stefani *et al*., 2015)) and the *lin-46* 5’UTR has been shown to prevent precocious accumulation of LIN-46 protein during development (Ilbay *et al*., 2021). To uncouple *lin-46* from LIN28, we mutated the single GGAG motif in the short endogenous *lin-46* 5’UTR to CTCC (*lin-46(xe370)*, henceforth referred to as *lin-46(++)*). To evaluate the levels of *lin-46* mRNA, we collected samples of synchronized animals hourly from 5 to 14 hours of development (i.e., from early/mid-L1 to early/mid-L2) and performed mRNA-seq. As shown in Fig. 5A, *lin-46* mRNA levels were upregulated to an intermediate level in *lin-46(++)* animals, with the characteristic increase at ∼8–9 hours observed in wild-type animals preserved (Fig. 5A, compare with Fig. 1D).

**Figure 5.**
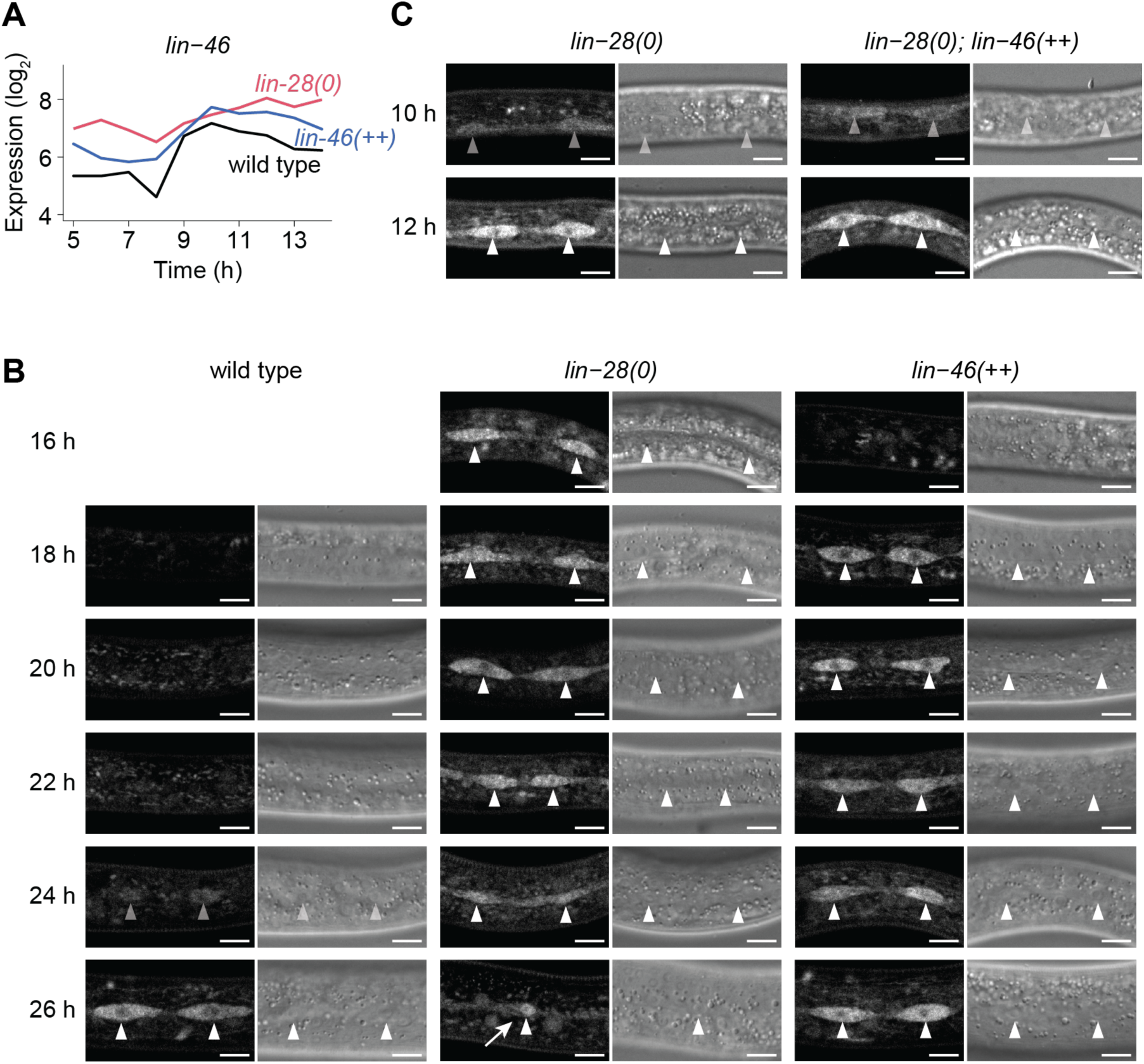
Mutation of the GGAG motif in *lin-46* 5’UTR recapitulates the precocious LIN-46 accumulation seen in *lin-28(0)* animals only partially. **A:** Levels of *lin-46* mRNA as determined by mRNA-seq of synchronized animals collected hourly from 5 to 14 h of development at 25°C. **B**: Confocal images of endogenously split-GFP-tagged LIN-46 in the epidermal seam cells of wild-type, *lin-46(++)* and *lin-28(0)* animals. Animals were grown at 25°C and examined at the indicated time points. Arrowheads indicate seam cells with detectable GFP signal. The arrow points to the predominantly nuclear accumulation of LIN-46 in *lin-28(0)* animals. Scale bars: 10 μm. **C**: Confocal images of endogenously split-GFP-tagged LIN-46 in the epidermal seam cells of *lin-28(0)* and *lin-28(0); lin-46(++)* animals. Animals were grown at 25°C and examined at the indicated time points. Arrowheads indicate seam cells with detectable GFP signal. Scale bars: 10 μm.

To determine whether this mild increase in *lin-46* mRNA levels also manifests at the protein level, we examined endogenous LIN-46 protein C-terminally tagged with a 4xsplit-GFP (*lin-46(xe347)*). In agreement with previously published observations using a different fluorophore (Ilbay *et al*., 2021), we detected LIN-46 in wild-type animals in the epidermal seam cells (Fig. 5B), as well as in the vulval precursor cells, tail and rectum (Fig. S4). Focusing on the seam cells, we observed LIN-46 accumulation starting in late L3 (24–26 h; Fig. 5B). In *lin-28(0)* mutant animals, LIN-46 accumulated two stages earlier, at late L1, from 12 hours onwards. This was true regardless of whether the *lin-46* 5’UTR was mutated (Fig. 5B, C). By contrast, and consistent with the RNA-seq data, in animals with the GGAG->CTCC mutation in the *lin-46* 5’UTR (*lin-46(xe351xe347)*), LIN-46 is detected at an intermediate time, in late L2 (from ∼18 hours onwards), one stage earlier than in wild-type but one stage later than in *lin-28(0)* (Fig. 5B). We noticed, but did not investigate further, that LIN-46 protein accumulates in the nucleus in *lin-28(0)* animals at ∼26 hours (Fig. 5B, white arrow).

**Figure S4.**
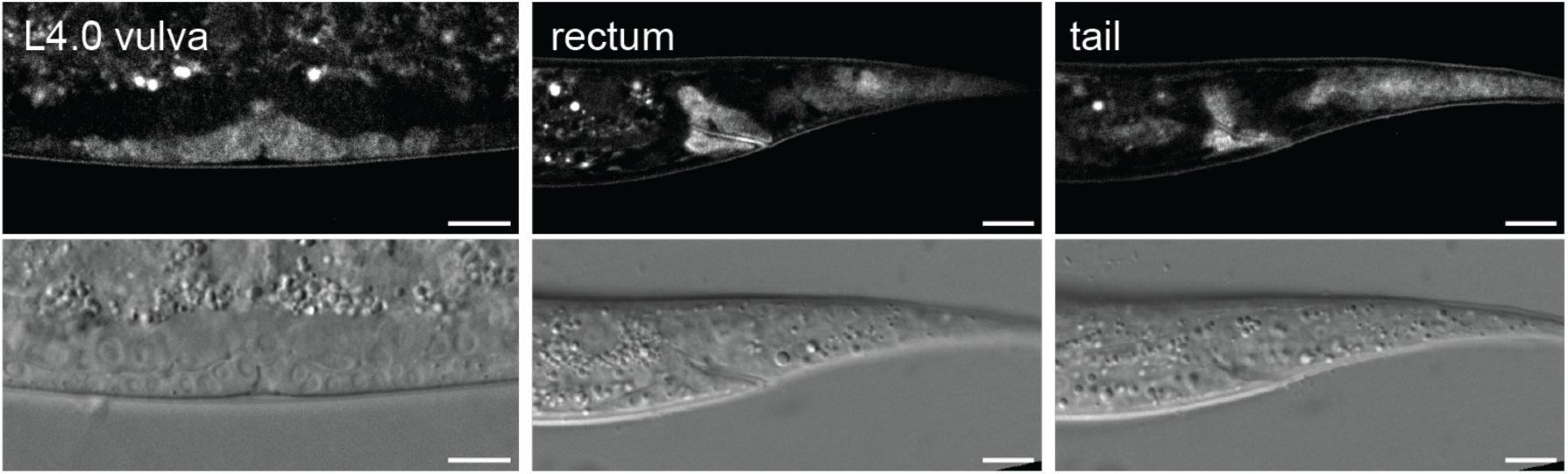
Spatial expression pattern of *lin-46* in wild-type L4 animals. Confocal images of endogenously split-GFP-tagged LIN-46 in the developing vulva (left), rectum (middle) and tail (right) of wild-type animals at 26 hours of development at 25°C. Scale bars: 10 μm.

Taken together, *lin-46* repression is only partially dependent on the single GGAG motif within its 5’UTR. The fact that some repression remains in *lin-46(++)* animals and is still LIN28-dependent suggests either that LIN28 has an additional binding site in the *lin-46* transcript, or that it additionally represses *lin-46* through an indirect mechanism.

### Enhanced expression of *let-7* and *lin-46* in two independent sensitized backgrounds emulates *lin-28* mutant phenotypes

Since the manipulation of putative LIN28 binding sites achieved only a partial decoupling of *lin-46* from LIN28-mediated regulation, insufficient to elicit heterochronic phenotypes (see below) and no decoupling of *let-7*, we performed further sufficiency testing in a sensitized genetic background. Specifically, LIN28 occurs in two isoforms, LIN28a and LIN28b, encoded in the same locus (Fig. 1A). The two proteins share the C-terminal part containing the RNA-binding domains but differ in their N-terminal sequences encoded by isoform-specific first exons. A deletion of the LIN28b*-*specific first exon in a *lin-28::flag::ha* background results in the complete loss of detectable LIN28b protein (Fig. S5A; *lin-28(xe344xe59)*, henceforth referred to as *lin-28(Δb)*) and an intermediate upregulation of both let-7 miRNA and *lin-46* mRNA throughout the examined time window (12 to 17 hours on food at 25°C; Fig. S5B). Despite these gene expression changes, the animals exhibit no signs of protruding vulva (Fig. 6A) or precocious alae synthesis (Fig. 6B). However, in the luciferase assay, *lin-28(Δb)* animals exhibit a fully penetrant 4M* phenotype (i.e., a defect in pharyngeal cuticle shedding during the fourth molt; see above; Fig. 6C, D).

**Figure S5.**
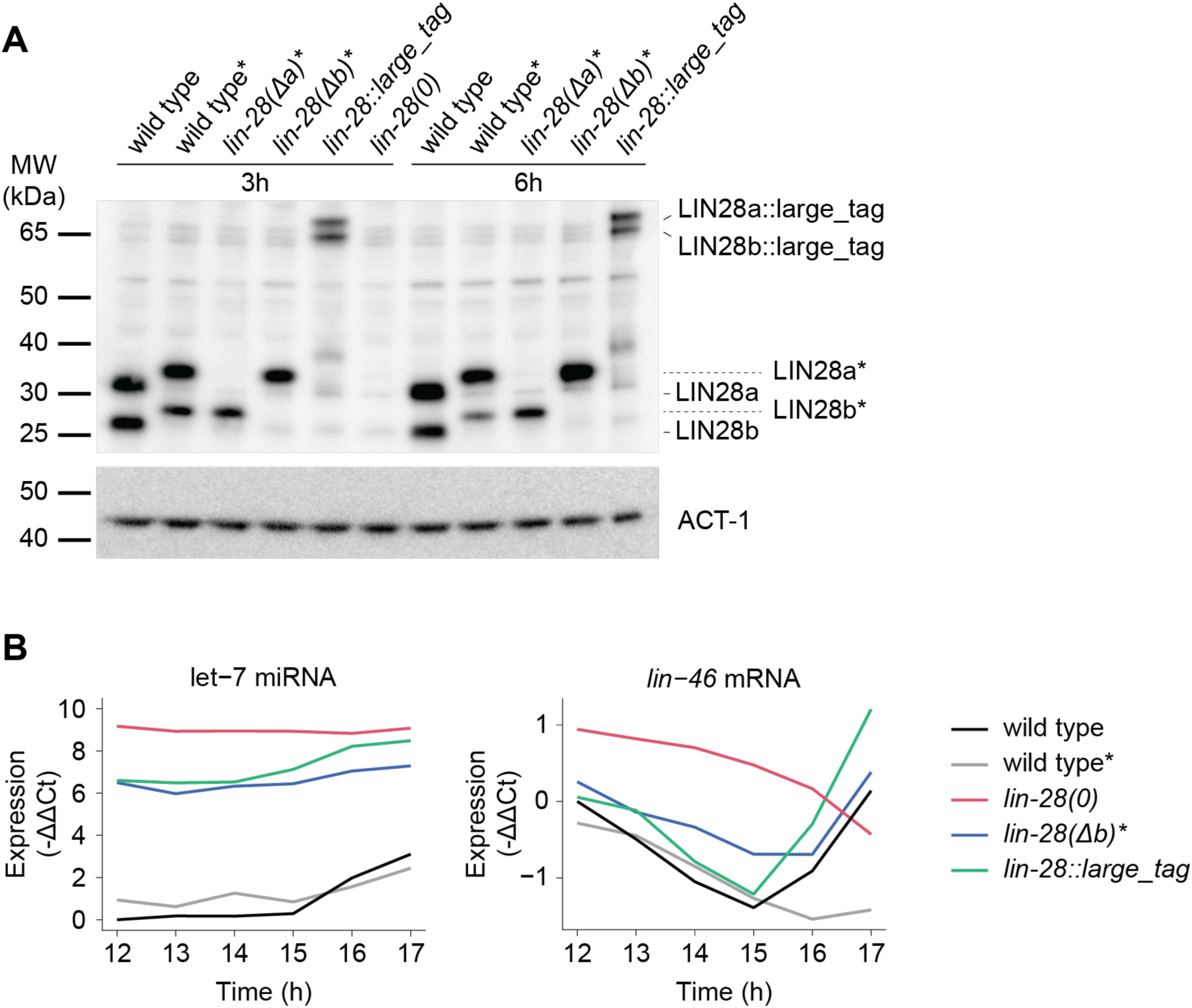
Generation of *lin-28(Δb)* isoform-specific mutant. **A**: Deletion of isoform-specific *lin-28* exons results in the loss of the respective protein isoform. Western blot showing the levels of both LIN28 isoforms, with size differences due to varying tag sizes in different genetic backgrounds. The LIN28a band is absent in *lin-28(Δa)* animals, whereas the LIN28b band is absent in *lin-28(Δb)* animals, validating the successful generation of isoform-specific mutants. As an antibody specificity control, *lin-28(0)* animals, which lack LIN28 protein, were included. Animals were grown at 25°C and collected at 3 and 6 h of development. Protein lysates were probed using anti-LIN28 and anti-actin antibodies. *: LIN28 is C-terminally FLAG::HA-tagged. **B**: Expression levels of mature let-7 miRNA (left) and *lin-46* mRNA (right) measured by RT-qPCR in animals grown at 25°C and collected hourly from 12 to 17 hours of development. Expression levels at each time point were first normalized to the levels of a reference gene (small nucleolar RNA sn2841 for let-7, actin mRNA for *lin-46*) and then to wild type at 12 h. *: LIN28 is C-terminally FLAG::HA-tagged.

**Figure 6.**
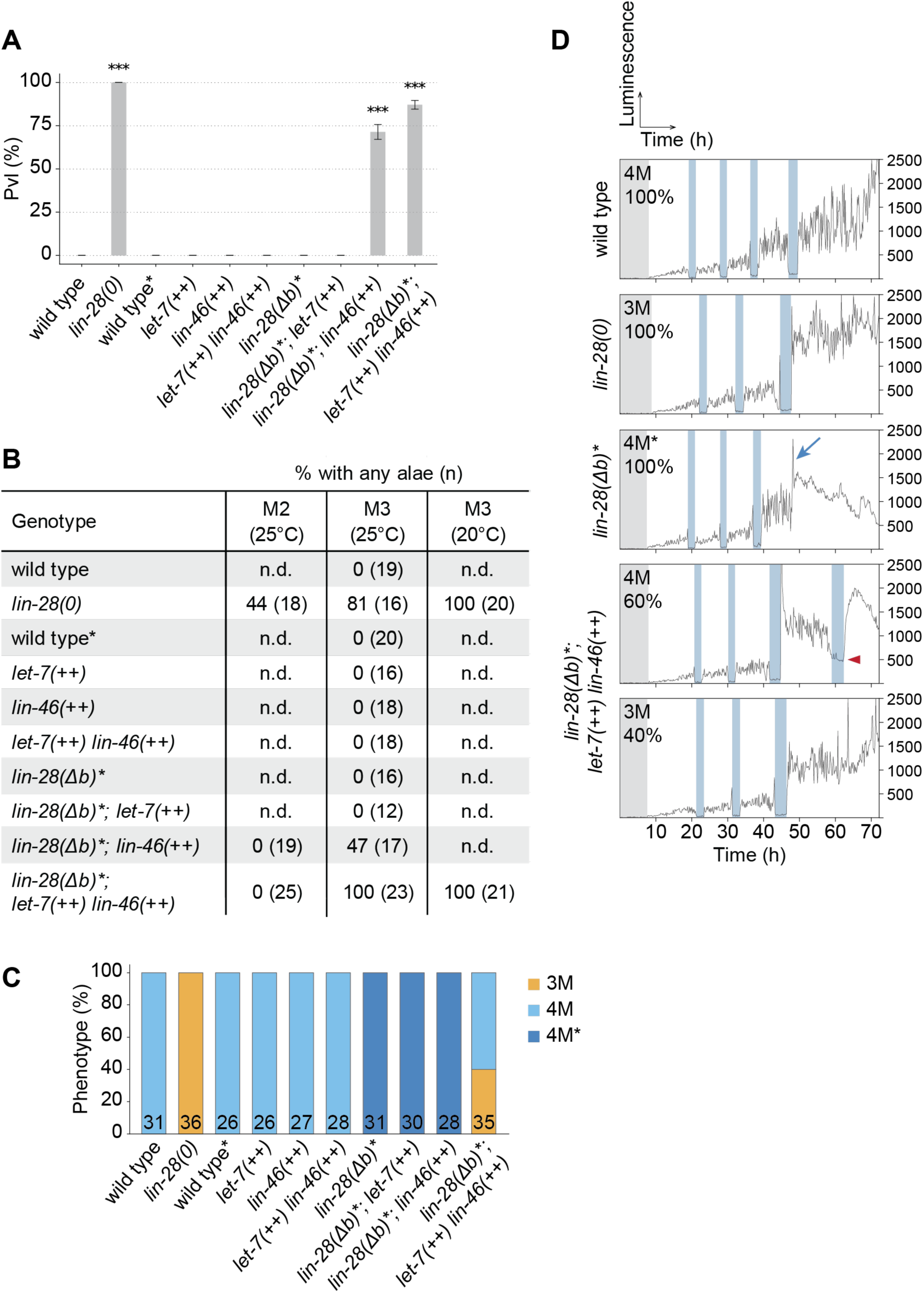
Enhanced expression of *let-7* and *lin-46* is sufficient to induce *lin-28* mutant-like phenotypes in *lin-28(Δb)* background. **A**: Quantification of the protruding vulva (Pvl) phenotype of adult animals of indicated genotypes. Animals were grown at 20°C and examined at 52 h of development, when they had reached adulthood. N = 4 technical replicates, with n > 230 animals scored per genotype. Error bars represent standard deviation. Significant differences to wild type were determined by a Dunnett’s test (\*\*\**p* < 0.001). **B**: Quantification of the presence of alae in animals of indicated genotypes. Molting animals were singled and examined shortly after they exited the respective molt as indicated by ecdysis. The wild type and *lin-28(0)* conditions are replotted from Fig. 3A for comparison. **C**: Quantification of the number of molts in animals of the indicated genotypes. The number at the bottom of each bar indicates the number of examined animals (n). **D**: Examples of luminescence intensity traces. Molt phenotypes are labeled as 3M for animals undergoing three molts, 4M for four molts, and 4M* for animals exhibiting an aberrant increase in luminescence intensity at the time of the expected fourth molt (indicated by a blue arrow). The two distinct outcomes of enhanced *let-7* and *lin-46* expression in the *lin-28(Δb)* background are illustrated. Note that in these animals, 4M involves a delayed and not properly executed fourth molt (4M), characterized by elevated luminescence intensity during this molt (arrowhead). Vertical blue bars indicate molts. The percentages indicate the penetrance of each phenotype (as shown in (**C**)). *: LIN28 is C-terminally FLAG::HA-tagged.

In the *lin-28(Δb)* background, we further enhanced the expression of *let-7* and/or *lin-46*, by introducing an additional *let-7* gene copy (hereafter *let-7(++)*), and by mutating the *lin-46* 5’UTR as in the *lin-46(++)* allele, respectively. Enhanced expression of *lin-46* in the *lin-28(Δb)* background resulted in partially penetrant protruding vulva (71%) and precocious alae (47%) phenotypes (*lin-28(Δb); lin-46(++)* in Fig. 6A, B). The penetrance of these phenotypes increased further when we additionally enhanced *let-7* expression (from 71% to 87% and from 47% to 100%, respectively; *lin-28(Δb); let-7(++) lin-46(++)* in Fig. 6A, B).

Upon individual enhanced expression of either *let-7* or *lin-46* in the *lin-28(Δb)* background, the animals still underwent the aberrant fourth molt characteristic of the 4M* phenotype (Fig. 6C). However, when we jointly enhanced the expression of both genes in this background, 40% of animals failed to undergo a fourth molt entirely, while the remaining animals underwent three proper molts followed by an incomplete drop in the luminescence signal, possibly reflecting an aberrant molt (Fig. 6C, D). We note that this latter molting pattern was distinct from the 4M* pattern, in which the signal steeply increases rather than decreases at the time of the expected fourth molt (Fig. 6D). In repeat experiments, we observed an even higher frequency of animals undergoing only three molts, namely 63% (n = 27) and 67% (n = 45).

These findings were qualitatively recapitulated using a second sensitized background, *lin-28::large_tag* (Fig. S6). These animals look overtly wild-type, but, as noted also by others (Nelson & Ambros, 2025), GFP-tagging subtly impairs LIN28 function, causing an increase in let-7 miRNA levels comparable to *lin-28(Δb)* animals (Fig. S5B).

**Figure S6.**
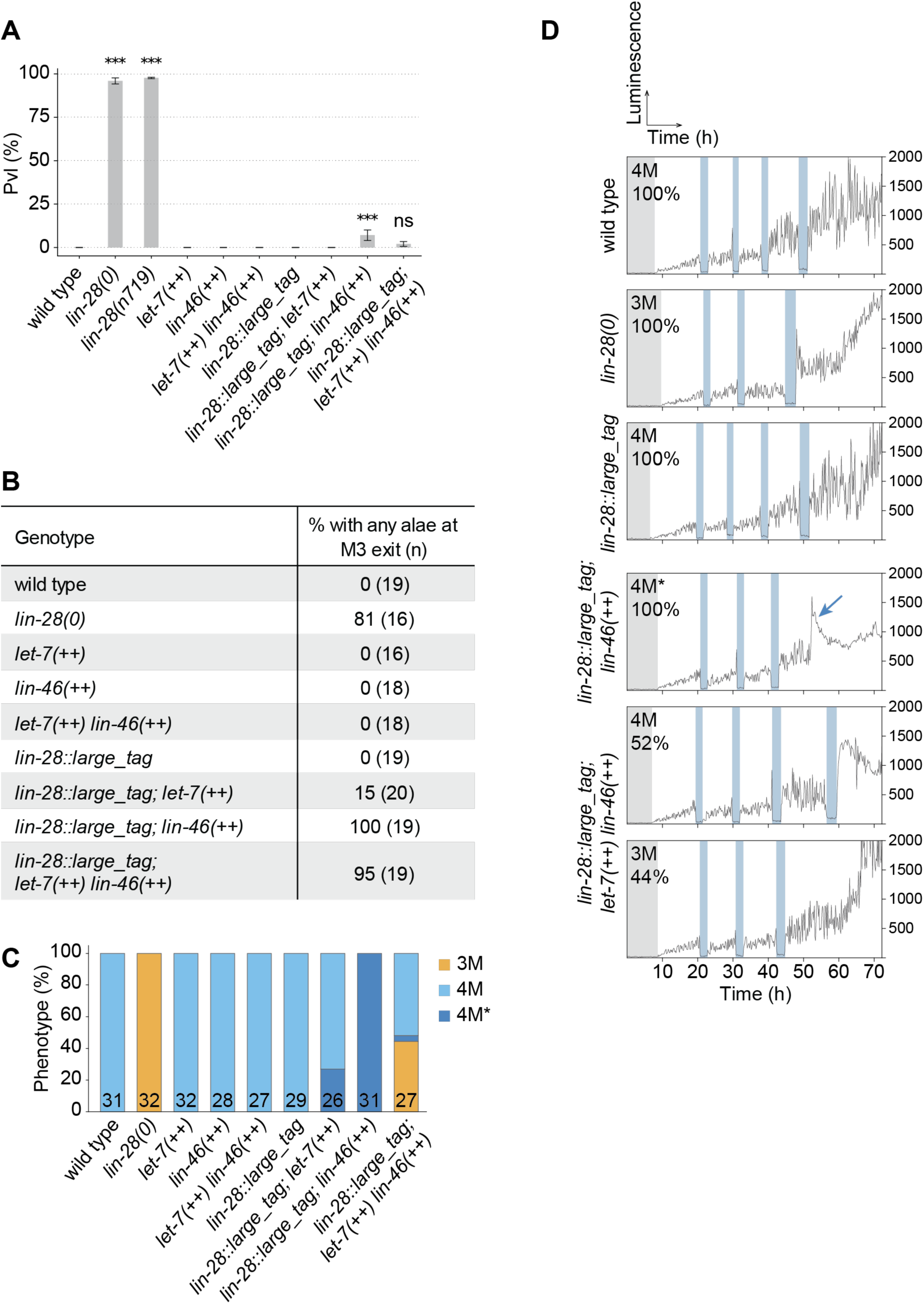
Enhanced expression of *let-7* and *lin-46* is sufficient to induce *lin-28* mutant-like phenotypes in *lin-28::large_tag* background. **A:** Quantification of the protruding vulva (Pvl) phenotype of adult animals of indicated genotypes. Animals were grown at 20°C and examined at 54 h of development. N = 4 technical replicates, with n > 170 animals scored per genotype. Error bars represent standard deviation. Significant differences to wild type were determined by a Dunnett’s test (ns: not significant; \*\*\**p* < 0.001). **B**: Quantification of the presence of alae in animals of indicated genotypes. Molting animals were singled and examined shortly after they exited the respective molt as indicated by ecdysis. The wild type and *lin-28(0)* conditions are replotted from Fig. 3A. **C**: Quantification of the number of molts in animals of the indicated genotypes. The number at the bottom of each bar indicates the number of examined animals (n). **D**: Examples of luminescence intensity traces. Molt phenotypes are labeled as 3M for animals undergoing three molts, 4M for four molts, and 4M* for animals exhibiting an aberrant increase in luminescence intensity at the time of the expected fourth molt (indicated by a blue arrow). The two distinct outcomes of enhanced expression of both *let-7* and *lin-46* in the *lin-28::large_tag* background are illustrated. Vertical blue bars indicate molts. The percentages indicate the penetrance of each phenotype (as shown in (**C**)).

### LIN28 controls not only the number of larval stages but also their duration

*C. elegans* larval development is temporally segmented such that cells assume specific identities characteristic of a given larval stage (Moss, 2007). Heterochronic genes were defined by their ability to specify these temporal fates, and their key role thus appears to be the maintenance of temporal order (Ambros & Horvitz, 1984). Other, clock-type mechanisms, involving oscillatory gene expression, are thought to set the tempo at which larval stages are executed (Olmedo *et al*, 2017; Tsiairis & Großhans, 2021). Surprisingly then, although *lin-28(0)* mutant animals execute only three instead of four larval stages, we found that the average duration from hatching to the exit from the final molt was comparable between wild-type and *lin-28(0)* animals (Fig. 7A): the final, third molt in *lin-28(0)* animals occurs only shortly before the fourth molt in wild-type (Fig. 7B). This is due to a substantial extension of larval stages in *lin-28(0)* animals, with the third larval stage being most strongly affected (Fig. 7C). This extended L3 stage phenotype depends on the expression of both *lin-46* and *let-7* (Fig. 7D, Fig. S7) and can be induced by their enhanced expression in sensitized genetic backgrounds with reduced LIN28 function (Fig. 7E, F).

**Figure 7.**
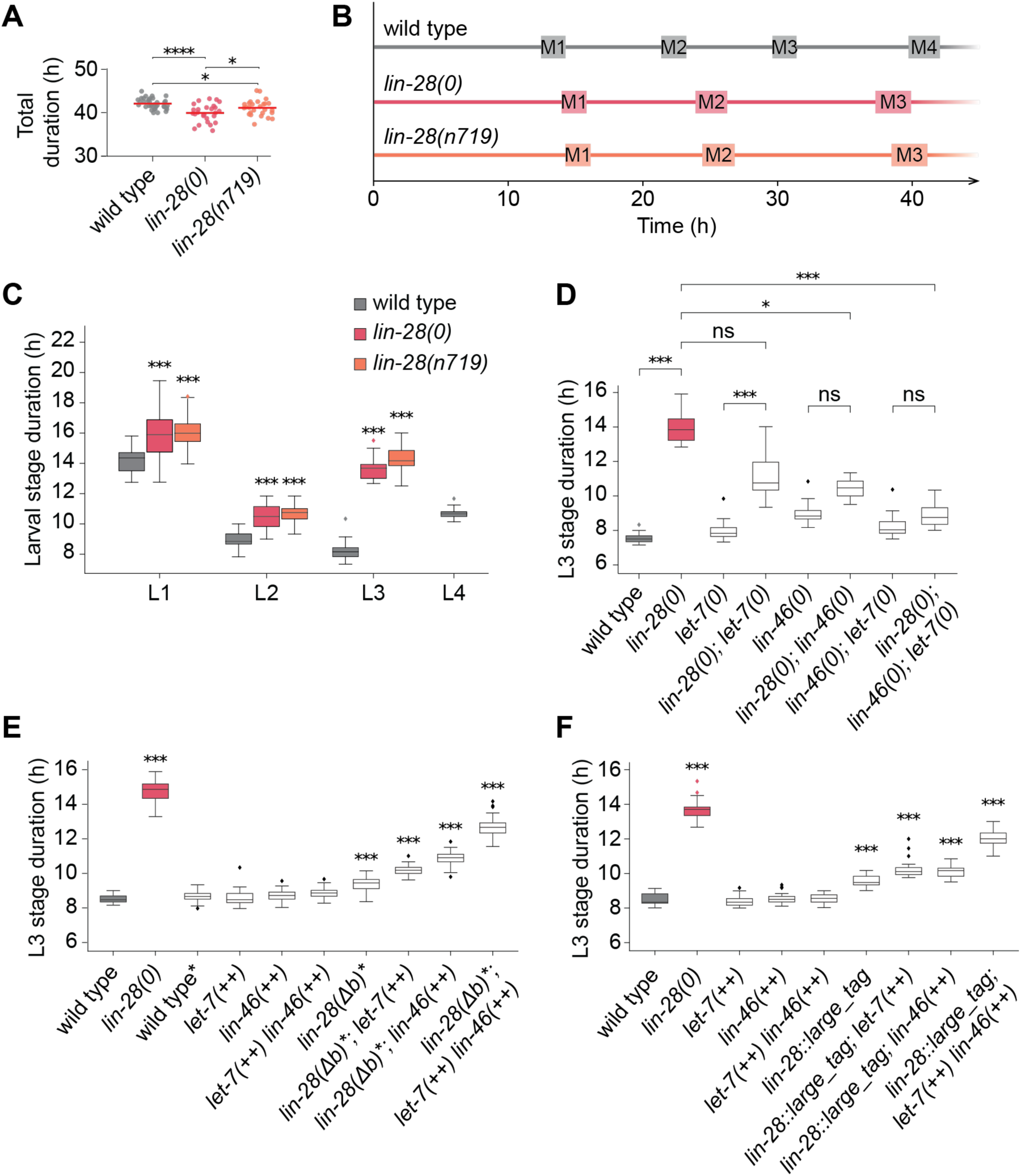
Lack of LIN28 results in extension of larval stage durations. **A**: Quantification of total duration of larval development from hatching until exit from the last executed molt. Number of animals n = 36, 27 and 27 for wild type, *lin-28(0*) and *lin-28(n719)*, respectively. The horizontal line represents the mean. Significant differences were determined by an unpaired two-sided *t*-test (\*\*\**p* < 0.001; \**p* < 0.05). **B**: Average duration of developmental stages in wild-type, *lin-28(0)* and *lin-28(n719)* animals as determined by the luciferase assay (same experiment as shown in (**A**)). Rectangles indicate the molts, horizontal lines represent the intermolts. **C**: Quantification of larval stage durations of wild-type, *lin-28(0)* and *lin-28(n719)* animals (same experiment as shown in (**A**)). The horizontal line represents the median, hinges extend to first and third quartiles, and the whiskers extend up to the most extreme value within 1.5*IQR (interquartile range) of the median. Significant differences relative to wild type for each larval stage are indicated. *P*-values were determined by a non-parametric Wilcoxon rank sum test. \*\*\**p* < 0.001. **D**: Quantification of the L3 stage duration of the same animals as in Fig. 4B. The horizontal line represents the median, hinges extend to first and third quartiles, and the whiskers extend up to the most extreme value within 1.5*IQR (interquartile range) of the median. Significant differences were determined by a Dunn’s test with Holm correction for multiple testing. The statistical significance of selected comparisons is indicated (\*\*\**p* < 0.001; \**p* < 0.05; ns, not significant). For statistical significance of all comparisons see Fig. S7. **E**: Quantification of the L3 stage duration of the same animals as in Fig. 6C, examining the effects of *lin-46* and *let-7* expression modulation in a *lin-28(Δb)* mutant background. The horizontal line represents the median, hinges extend to first and third quartiles, and the whiskers extend up to the most extreme value within 1.5*IQR (interquartile range) of the median. Significant differences to wild type were determined by a Dunn’s test with Holm correction for multiple testing (\*\*\**p* < 0.001; no label, not significant). **F**: Quantification of the L3 stage duration of the same animals as in Fig. S6C, examining the effects of *lin-46* and *let-7* expression modulation in a *lin-28::large_tag* background. The horizontal line represents the median, hinges extend to first and third quartiles, and the whiskers extend up to the most extreme value within 1.5*IQR (interquartile range) of the median. Significant differences to wild type were determined by a Dunn’s test with Holm correction for multiple testing (\*\*\**p* < 0.001; no label, not significant). *: LIN28 is C-terminally FLAG::HA-tagged.

**Figure S7.**
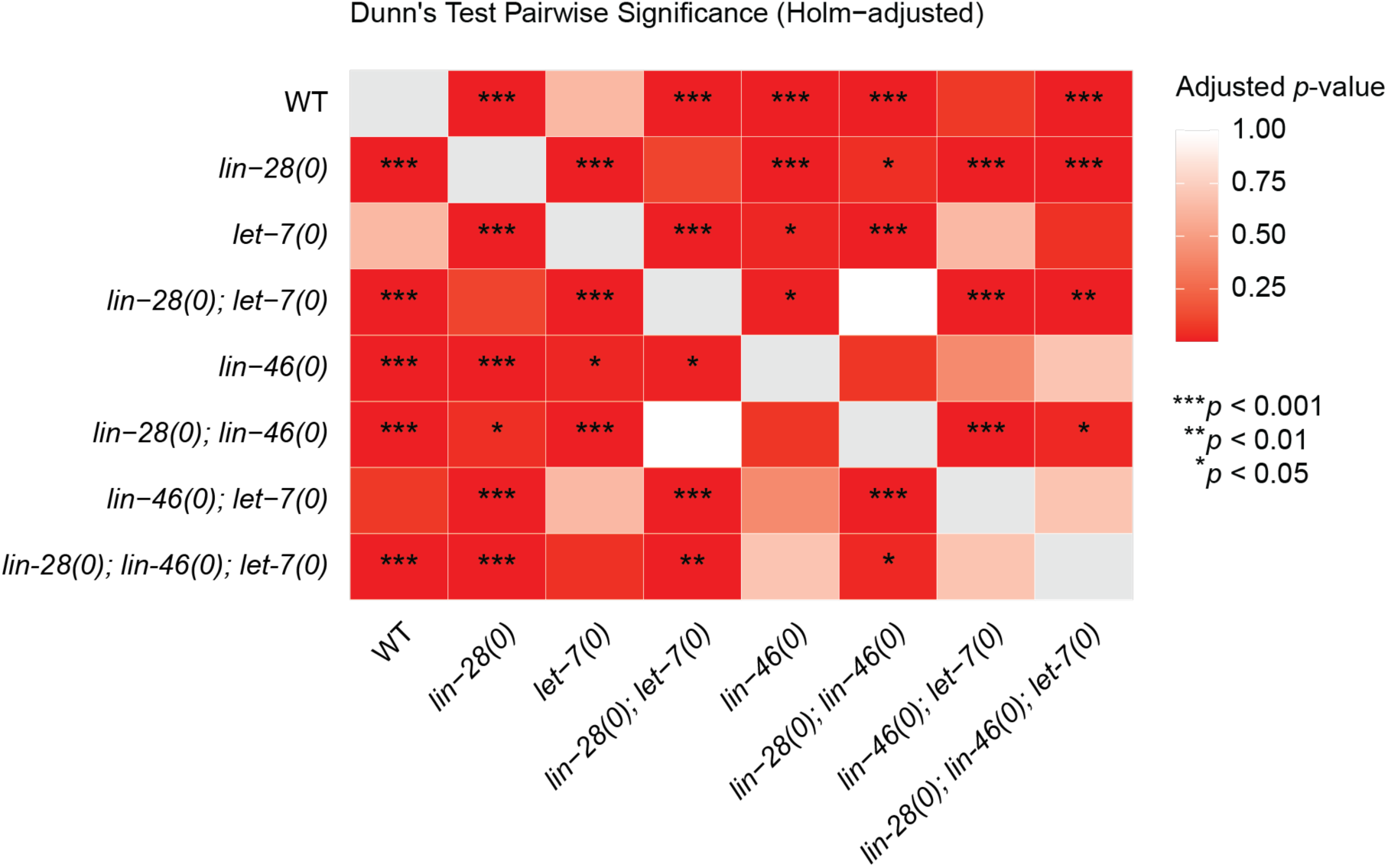
Statistical significance of pairwise L3 duration comparisons. Relates to Fig. 7D. A heatmap showing adjusted *p*-values from pairwise non-parametric comparisons between L3 stage durations of tested genotypes determined by a Dunn’s test with Holm correction for multiple testing (\*\*\**p* < 0.001; \*\**p* < 0.01; \**p* < 0.05; no label, not significant).

In wild-type animals, the expression of thousands of genes is rhythmic, peaking once per larval stage (Fig. 8A, B and (Hendriks *et al*., 2014; Meeuse *et al*., 2020)). Consistent with the reduced number of molts accompanied by the gradually increasing extension of larval stages, oscillating genes peaked only three times in *lin-28(0)* animals over the course of the experiment (Fig. 8A, B), with an increased period (Fig. 8C, D).

**Figure 8.**
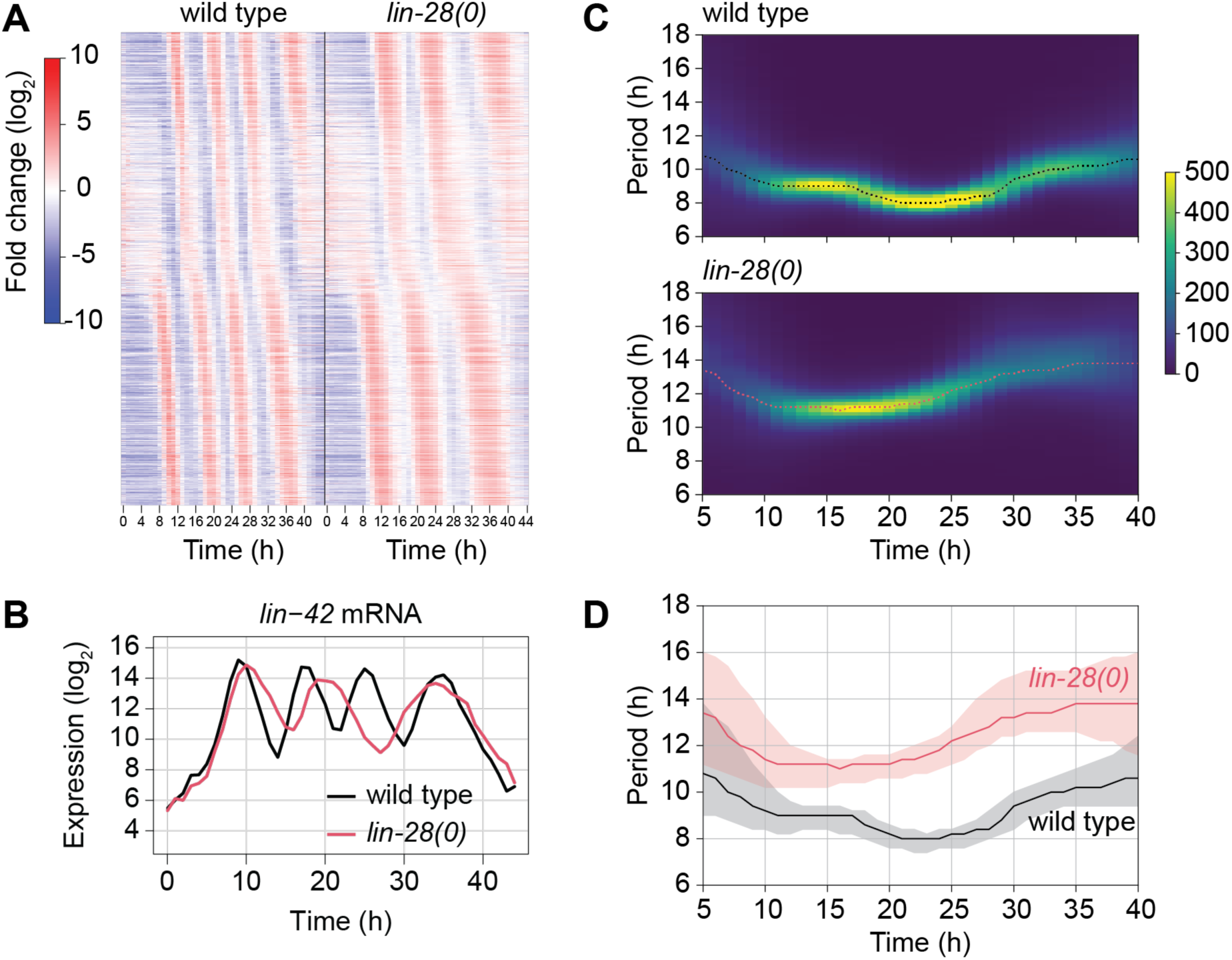
Oscillatory gene expression reflects the perturbed molting cycle in *lin-28(0)* animals. **A**: Heatmap showing mean-normalized log_2_ expression levels of oscillating genes (n = 2,004; as defined in (Meeuse *et al*., 2020), with log_2_ amplitude ≥ 1 in that study) over time in wild-type (left) and *lin-28(0)* animals (right). Genes are ordered according to the published peak phase. **B**: Expression profile of a representative oscillating gene, *lin-42*, across development in wild-type and *lin-28(0)* animals. **C**: Distribution of instantaneous periods of all oscillating genes (n=3,680) in wild-type and *lin-28(0)* animals. Color bar represents number of genes per bin at each timepoint. Average period is shown as a dashed line. The instantaneous period for each gene was computed using the Hilbert Transform on the data after processing it with a Butterworth filter. **D**: Comparison between the average periods replotted from (**C**). The shadowed area corresponds to the values between the 20^th^ and 80^th^ percentiles, from the distributions shown in (**C**).

Taken together, our findings show that LIN28 controls not only the identity of larval stages but also their duration. Whether this extends to other heterochronic genes and through what molecular mechanism remains to be elucidated.

## Discussion

First discovered as a regulator of temporal patterning in *C. elegans*, LIN28 also plays various roles in humans, from acting as a pluripotency factor to controlling the onset of puberty. In various contexts, LIN28 proteins bind hundreds of transcripts, fitting the stereotype of RNA-binding proteins as promiscuous binders. However, here we have shown that in *C. elegans*, regulation of only two targets, *let-7* and *lin-46*, is sufficient to explain the developmental function of LIN28. Previous studies had shown that individual mutation of *let-7* or *lin-46* could partially suppress *lin-28* mutant precocious heterochronic phenotypes (Pepper *et al*., 2004; Reinhart *et al*., 2000; Slack *et al*., 2000; Vadla *et al*., 2012). Yet, the resulting double mutant animals did not recapitulate the retarded heterochronic phenotypes of *let-7* or *lin-46* loss, respectively, but instead exhibited an “in-between” phenotype. Such mutual suppression supported the existence of additional targets. We demonstrate here that *lin-28; lin-46; let-7* triple null mutant animals recapitulate *lin-46; let-7* double null mutant phenotypes. In other words, mutation of these two target genes is fully epistatic to mutation of *lin-28*.

While *lin-46* and *let-7* are both required for *lin-28* precocious phenotypes related to molting and cuticle synthesis, *lin-46* is the dominant factor for the protruding vulva phenotype and, not re-examined by us, the skipping of the L2-specific symmetric seam cell division, which is suppressed by loss of *lin-46* (Pepper *et al*., 2004) but not of *let-7* (Vadla *et al*., 2012). This overall distinction agrees with the earlier proposal that LIN28’s function in regulating cell proliferation precedes later activities (Vadla *et al*., 2012). Furthermore, this may also explain why mutation of the *lin-46* 5’UTR, which advances LIN-46 accumulation by only one stage as opposed to two in *lin-28* mutant animals, is insufficient for a Pvl phenotype. (We note that small deletions in the *lin-46* 5’UTR have been reported to induce a Pvl phenotype (Ilbay *et al*., 2021), but our data suggest a contribution from additional factors in that strain’s background (see Methods).)

Interestingly, these findings also indicate that dysregulation of LIN-46 rather than let-7 appears to be the main driver of *lin-28* mutant phenotypes, despite the extensive phylogenetic conservation of let-7 regulation by LIN28. In other words, early larval development is more robust to elevated levels of let-7 than LIN-46. In contrast, not having enough let-7 later in development is lethal, whereas LIN-46 loss is well tolerated – possibly due to functional redundancies between LIN-46 and the let-7 sister miRNAs miR-48, miR-84 and miR-241 (Ilbay & Ambros, 2019).

Despite the conservation of *let-7* as a LIN28 target and the mechanism of its regulation, its mode of recognition in *C. elegans* remains to be elucidated. Whereas mammalian LIN28 recognizes a GGAG motif in the loop of *pri-* and *pre-let-7* stem-loop structures (Loughlin *et al*., 2011; Nam *et al*., 2011; Ustianenko *et al*., 2018), *C. elegans let-7* lacks this motif. CLIP experiments instead suggested binding to the *pri-let-7* outside the stem-loop structure (Stefani *et al*., 2015), yet when we mutated the four implicated GGAG motifs, only minor, statistically insignificant changes in mature let-7 levels resulted, in contrast to the >50-fold upregulation observed upon LIN28 loss at the same early developmental stage. Similarly, mutation of the GGAG motif in the *lin-46* 5’UTR caused much weaker derepression than LIN28 loss. Given that binding of LIN28 to *lin-46* was in fact detected mostly outside the 5’UTR (Stefani *et al*., 2015), it seems possible that additional, redundant binding sites exist, and this may also be true for *let-7*. These binding sites might also be distinct from the canonical GGAG motif, as LIN28’s cold-shock domain is thought to promote transcript binding through a distinct GAU motif (Loughlin *et al*., 2011; Nam *et al*., 2011; Ustianenko *et al*., 2018). As this motif is likewise absent from both the apical part of the *C. elegans pri-let-7* stem-loop and the *lin-46* 5’UTR, the mode of *C. elegans* LIN28 binding to its targets remains to be elaborated.

It had previously been speculated that LIN28 regulates *lin-46* at the level of translation (Ilbay *et al*., 2021). Nonetheless, we reproducibly observed increases in *lin-46* mRNA levels in both animals with mutated *lin-28* or mutated LIN28 bindings site in the *lin-46* 5’UTR. We suspect that the dynamic changes in lin-46 mRNA levels in wild-type animals might have obscured the effect of *lin-28* loss in the earlier work. It also remains possible that, similar to the situation with miRNAs, *lin-46* silencing involves both degradation and translational repression.

Our data do not formally rule out the possibility that LIN28 alters the activity of additional genes by translation or other mechanisms not captured by RNA-seq. Yet, if such regulation, not examined by us, exists, it appears largely inconsequential for the developmental functions that we have investigated here. Whether a similar scenario applies to mammalian LIN28 remains to be determined: regulatory roles were proposed for dozens of its bound transcripts (Maklad et al., 2023), but experimental validation of the physiological relevance is limited for most of these interactions.

In addition to LIN28, several more RNA binding proteins and miRNAs operate in the heterochronic pathway. Surprisingly, although miRNAs and RBPs are often considered to act inherently promiscuously, for the heterochronic pathway, a picture of great functional specificity is emerging: key factors, i.e., the lin-4 and let-7 miRNAs, the LIN-41 RBP and now also LIN28 all act through few (≤4) targets (Aeschimann *et al*, 2017; Aeschimann *et al*, 2019; Ecsedi *et al*, 2015; Moss *et al*., 1997; Wightman *et al*, 1993). This begs the question whether this is a unique feature of these particular factors and/or this particular pathway, or whether the possibility that RBPs and miRNAs act more specifically should be entertained also in other contexts. At least for miRNAs, more examples of uniquely relevant targets exist (Dorsett *et al*, 2008; Drexel *et al*, 2016; Johnston & Hobert, 2003; Lu *et al*, 2015; Teng *et al*, 2008). We do emphasize that greater specificity does not exclude the possibility of distinct targets in different contexts, and we envision a continuum, where some miRNAs (and RBPs) may have highly specific functions and others work more by a network-type mechanism. Nonetheless, pervasive RNA binding should not be taken at face value as an indicator of network activity, as high-throughput RBP-RNA binding data appear generally limited in their predictive power for functional interactions, likely due to a combination of technical limitations of available protocols and the capture of non-productive interactions (Chen *et al*, 2019; Vieira-Vieira & Selbach, 2021; Welte *et al*, 2023).

Finally, although *C. elegans* heterochronic genes have thus far been viewed specifically as temporal patterning genes that determine the temporal identity of cells, and thereby the order of developmental events, we show here that LIN28 loss additionally causes a strong decrease in developmental tempo, most pronounced in L3. The developmental slowing occurs rather uniformly across animals and rhythmic gene expression patterns remain robustly detectable at the population level, pointing to a specific effect rather than a mere deterioration of developmental robustness – a notion further supported by the fact that the additional loss of *let-7* and *lin-46* reverts developmental tempo to wild-type levels.

The clock that controls developmental tempo was previously shown to slow down during the L4 stage, while maintaining a constant amplitude of periodic gene expression, before ultimately settling into a stable, non-oscillatory regime as animals reach adulthood (Meeuse *et al*., 2020). Hence, a parsimonious explanation of the reduced developmental tempo in *lin-28* mutant animals is that loss of LIN28 pushes the system precociously towards the bifurcation point that separates the oscillatory from the non-oscillatory regime, without immediately crossing it. Clearly, the heterochronic pathway and the developmental clock are interconnected in numerous and complex ways that await further dissection in the future.

### Limitations of the study

The RIP-seq method that we used to identify mRNAs bound by LIN28 may also identify non-physiological binding partners, that emerged from post-lysis associations. However, since the overlap to previously published CLIP-seq data (Stefani *et al*., 2015) was substantial and significant, and *lin-46,* the only dysregulated gene, was also LIN28-bound, false positive binding events are not expected to impact the key conclusions.

As we were unable to effectively uncouple *let-7* and *lin-46* from regulation by LIN28, we had to rely on a sensitized background for sufficiency testing. Although similar results were obtained with two different sensitized backgrounds, it remains possible that this sensitization involves upregulation of additional LIN28 targets not captured by mRNA-seq. Future work, achieving complete decoupling of the two targets from LIN28-mediated repression without perturbation of *lin-28* itself, will be able to rule in or out this possibility. Irrespectively, when viewed together with the analysis of the suppression analysis, it is evident that *lin-46* and *let-7* are the functionally essential LIN28 targets.

## Material and Methods

### C. elegans handling

The wild-type strain used in this study was Bristol N2. Transgenic strains used in this study are listed in Table S1. To synchronize animals, arrested L1 stage larvae were obtained by extracting embryos from gravid adults using a bleaching solution (30% (v/v) sodium hypochlorite 4-6% reagent (Sigma-Aldrich; 419550010), 750 mM KOH) and hatching overnight in the absence of food, at room temperature in M9 buffer (42 mM Na_2_HPO_4_, 22 mM KH_2_PO_4_, 86 mM NaCl, 1 mM MgSO_4_). Experiments were performed at 25°C unless explicitly stated otherwise.

### Generation of transgenic CRISPR strains

Alleles generated in this study are listed in Table 1. Sequences of oligos used for CRISPR strain generation are listed in Table S2.

**Table 1.**
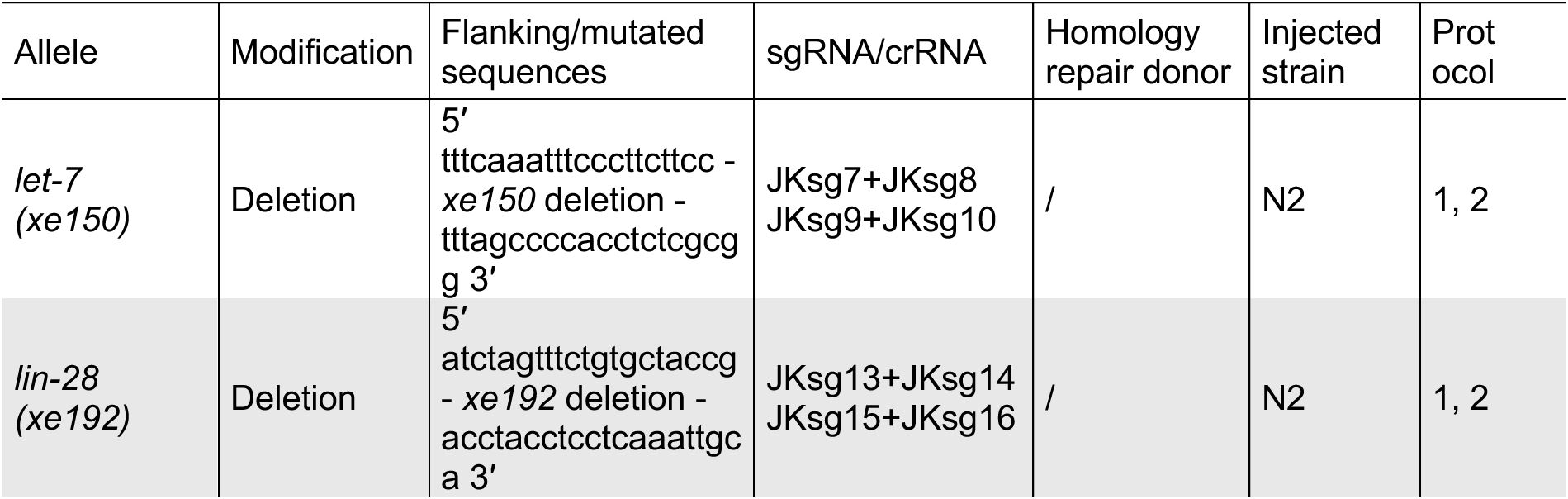
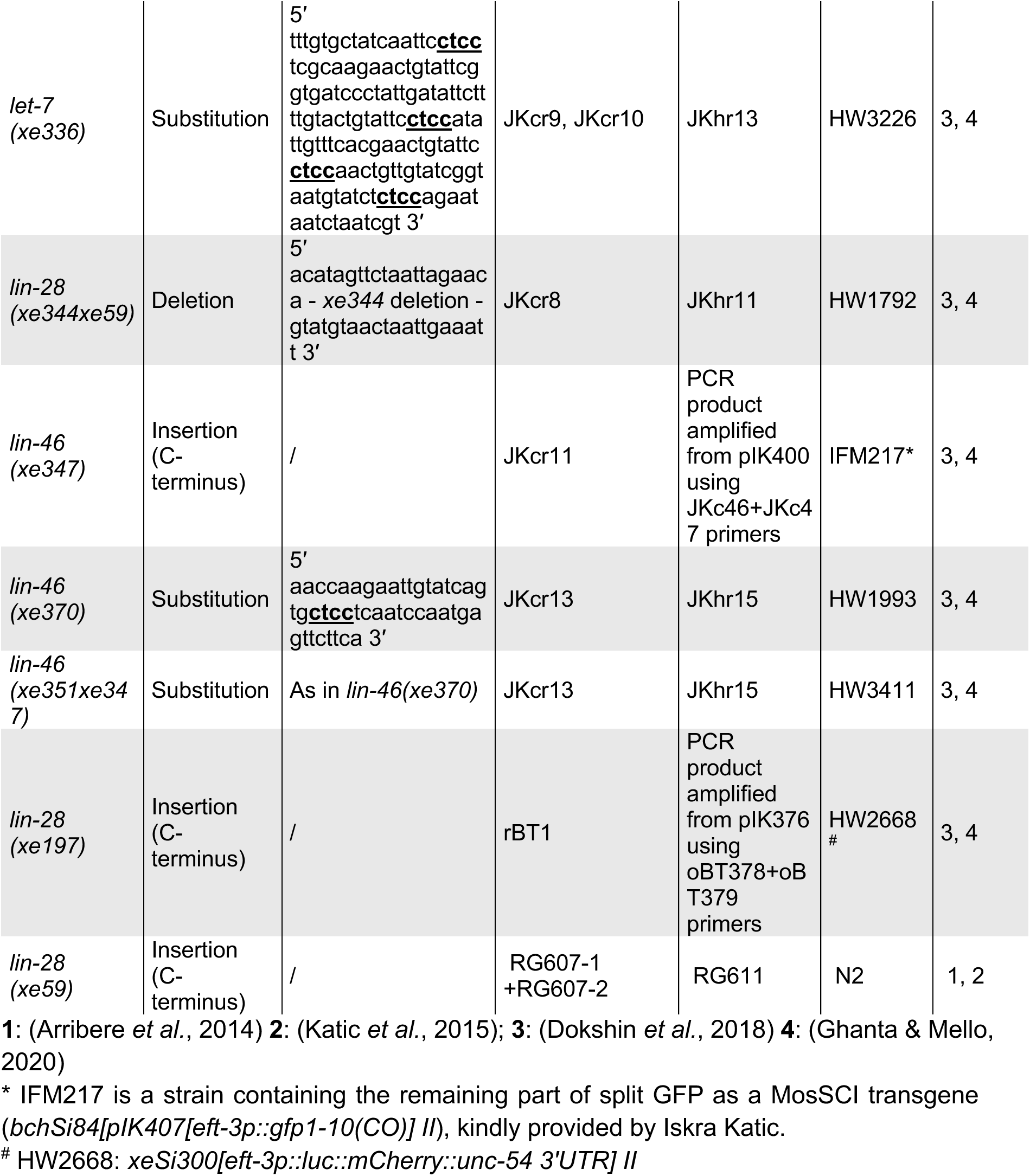
Alleles generated in this work.

The *let-7(xe150)* and *lin-28(xe192)* alleles were generated following an adapted “*dpy-10* conversion protocol” (Arribere *et al*, 2014; Katic *et al*, 2015). sgRNAs were cloned into a *Not*I-digested pIK198 plasmid using Gibson cloning. The injection mix contained: 50 ng/μl pIK155 (Cas9 expression vector), 100 ng/μl pIK208 (sgRNA expression vector targeting *dpy-10*), 20 ng/μl AF-ZF-827 (homology repair oligo for *dpy-10*) and 100 ng/μl of each pIK198 with a cloned sgRNA targeting the gene of interest.

The *let-7(xe336)*, *lin-28(xe344xe59)*, *lin-46(xe347)*, *lin-46(xe351xe347)*, *lin-46(xe370)* and *lin-28(xe197)* alleles were generated following published protocols (Dokshin *et al*, 2018; Ghanta & Mello, 2020). crRNAs targeting the genes of interest were ordered from IDT (Alt-R® CRISPR-Cas9 crRNA, 2 nmol, standard desalting). A pre-injection mix with 0.5 μl Alt-R *S.p.* Cas9 Nuclease V3 (10 μg/μl; IDT, 1081058), 5 μl tracrRNA (0.4 μg/μl; IDT) and 2.8 μl crRNA (0.4 μg/μl) (or 1.4 μl of each crRNA if two were used) was incubated for 15 minutes at 37°C before adding 2 μl pGH8 (100 ng/μl), 2 μl pRF4 (400 ng/μl) and either 500 ng of melted dsDNA donor or 2.2 μl single-stranded oligo donor (1 μg/μl; ordered as 4 nmole Ultramer DNA Oligo Standard Desalting from IDT).

To test whether LIN28-mediated *lin-46* repression depends on the single GGAG motif present in *lin-46* 5’UTR, we changed it from GGAG to CTCC, generating the *lin-46(xe370)* allele. *lin-46(xe370)* animals did not exhibit a protruding phenotype (Results) unlike other *lin-46* 5’UTR mutant lines previously described (Ilbay *et al*., 2021). To address whether these differences reflected differences in the mutation or strain background, as the previously reported lines also contained a multicopy array integrated in the same chromosome as *lin-46 (maIs105 [col-19p::gfp] V)*, we sought to outcross the *maIs105* array from the *lin-46(ma467)* allele, a 12 nucleotide long deletion removing the first two G’s of the GGAG motif and ten nucleotides upstream (Ilbay *et al*., 2021). Despite screening over 300 F2 progeny from three separate outcross attempts, we did not detect any recombination events between the two loci, suggesting either that their genetic distance is small, or that crossover is suppressed. As an alternative approach, we introduced deletion identical to *ma467* in either wild-type or *lin-46::3xflag::gfp11* backgrounds using CRISPR/Cas9. This did not cause precocious LIN-46 accumulation or a protruding vulva phenotype. We speculate that the phenotypes observed by (Ilbay *et al*., 2021) might be synthetically induced by a genetic interaction of the *lin-46* 5’UTR deletions with the integrated *maIs105* array or reflect CRISPR off-target effects.

### Experiments to quantify gene expression

To quantify gene expression using sequencing techniques or RT-qPCR, animals were sampled either hourly over a developmental timecourse (denoted as “TC” in Table 2), or only at selected time points (denoted as “Exp” in Table 2). Samples were collected as described below. Prior to RNA extraction, animals were lysed by five cycles of thawing (42°C) and snap-freezing in liquid nitrogen. RNA was extracted either using Direct-zol RNA Microprep kit (Zymo Research; R2062; DNase treatment included) according to the manufacturer’s protocol or performing phenol-chloroform extraction according to the Tri Reagent (Molecular Research Center, TR-118) manufacturer’s protocol followed by DNase treatment (RNase-Free DNase I kit (#25710 NORGEN); Single Cell RNA Purification Kit (#51800 NORGEN)).

**Table 2.**
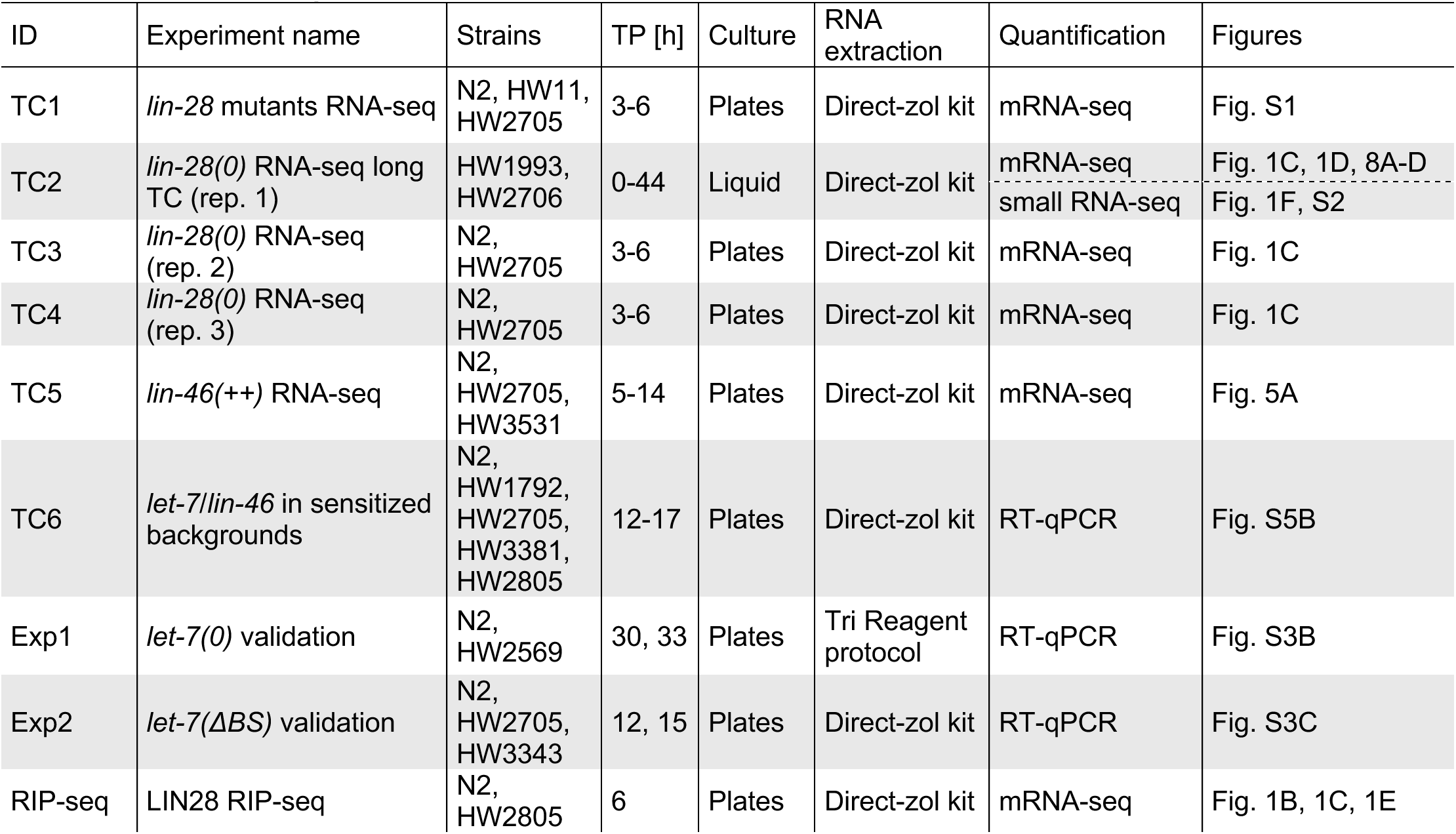
Overview of experiments.

#### Sample collection: animals grown on plates

Synchronized L1 animals were plated on 2% NGM plates seeded with OP50 *E. coli* bacteria and grown at 25°C. At the time of sample collection (Table 2), animals were washed off the plates with M9 buffer, washed twice with M9 buffer, resuspended in TRI Reagent and snap-frozen in liquid nitrogen.

#### Sample collection: animals grown in liquid culture

For the full developmental timecourse (TC2), synchronized L1s were grown in liquid culture (S-Basal supplemented with OP50; OD600 = 2.7; 1000 animals/ml) at 25°C shaking at 180 rpm. Animals were sampled hourly, washed three times with M9 buffer, resuspended in TRI Reagent and snap-frozen in liquid nitrogen.

#### TaqMan RT-qPCR: Quantification of mature let-7 miRNA

To quantify the levels of mature let-7 miRNA, 20 ng of total RNA was reverse transcribed using TaqMan MicroRNA Reverse Transcription Kit (Thermo Fisher Scientific; 4366596) and the following TaqMan MicroRNA Assays (Thermo Fisher Scientific): mature let-7 (ID 000377), sn2841 (reference gene; ID 001759). The reverse transcription (RT) reaction of total volume 15 μl contained 0.75 μl of each RT primer (for let-7 and sn2841) and was incubated at 16°C for 30 min, 42°C for 30 min and 85°C for 5 min. The resulting cDNA was used as template for qPCR. qPCR was performed either in Step one Realtime PCR (Thermo Fisher Scientific) or Roche LightCycler 480 machine depending on the number of reactions. A qPCR reaction contained TaqMan 2x Universal PCR Master Mix, no Amp Erase (Thermo Fisher Scientific; 4324018) and the above-listed TaqMan MicroRNA Assays. The PCR conditions were initiated with 10 min incubation at 95°C, followed by 40 cycles of 95°C for 15 s and 60°C for 60 s. Average Ct values were obtained from three technical replicates for each condition. Delta Ct (dCt) was calculated by subtracting Ct values of sn2841 from let-7 miRNA Ct values. Expression levels were normalized to a reference sample and plotted as -ddCt, where -ddCt = -(dCt – dCt_reference_). The N2 sample of each biological replicate (Fig. S3B, S3C) and the N2 12 h sample (Fig. S5B) were used as reference samples. *P*-values were determined by a paired two-sided *t*-test.

#### RT-qPCR: Quantification of *lin-46* mRNA

To quantify the levels of *lin-46* mRNA, 300 ng of total RNA was reverse transcribed to cDNA using the ImProm-II™ Reverse Transcription System (Promega; A3800) and oligo(dT) primers according to the manufacturer’s protocol. cDNA was diluted 1:400 to detect actin or used undiluted to detect *lin-46*. RT-qPCR reactions were performed using PowerUp™ SYBR™ Green Master Mix (Applied Biosystems; A25776) in Roche LightCycler 480 machine. Average Ct values were obtained from three technical replicates for each condition. Delta Ct (dCt) was calculated by subtracting Ct values of actin from *lin-46* Ct values. Expression levels were normalized to the N2 12 h sample (= reference sample) and plotted as -ddCt, where -ddCt = -(dCt – dCt_reference_) (Fig. S5B).

#### mRNA-seq

Sequencing was performed according to the selected protocol (Table 3). RNA integrity was validated using either RNA Nano Chip with Agilent 2100 Bioanalyzer or RNA ScreenTape with Agilent TapeStation. The quality control of the sequencing libraries was performed using High Sensitivity DNA Assay with Agilent 2100 Bioanalyzer.

**Table 3.** Overview of protocols used for mRNA-seq.

| ID | Sequencing protocol |
| --- | --- |
| TC1 | Paired-end Illumina NovaSeq 6000 |
| TC2 | Single-end Illumina HiSeq 2500 |
| TC3,4 | Paired-end Illumina NovaSeq 6000 |
| TC5 | Paired-end Illumina NovaSeq 6000 |

#### mRNA-seq: DEG in *lin-28(0)* vs. wild type

Sequencing reads were mapped to the *C. elegans* reference genome (ce11) with the R package QuasR using the spliced alignment algorithm HISAT2 (Gaidatzis *et al*, 2015). Protein-coding exons were defined based on the exon annotation file from WormBase (WS270) and read mappings to the corresponding genes were counted using qCount function from the QuasR package. To determine differentially expressed genes in *lin-28(0)* animals relative to wild type, three biological replicates of a developmental time window of 3 to 6 h were analyzed (TC2, TC3 and TC4; Fig. 1C). Read counts for each sample were pooled across the four time points (3 h, 4 h, 5 h and 6 h) and then normalized to the mean mapped library size, where ‘library size’ refers to the total number of mapped reads of the pooled libraries. The normalized counts were log₂-transformed with a pseudocount of 8. Differentially expressed genes were identified using the R package edgeR, applying batch correction for biological replicates. Genes with low expression were filtered out using filterByExpr. Quasi-likelihood F-tests were performed to assess differential expression, and genes with a false discovery rate (FDR) < 0.01 were considered significantly differentially expressed.

#### mRNA-seq: Spectral analysis

The dynamics of each previsously defined oscillating gene (Meeuse *et al*., 2020) (n = 3680), was processed using the Python (3.11) implementation of the Butterworth filter in the Scipy package (version 1.16.0, functions *butter, sosfiltfilt* and *hilbert* from the *signal* module). A first order bandpass filter was used, with critical frequencies of (0.07, 0.14) for the control and (0.06, 0.12) for the mutant strain. These frequencies were chosen empirically to span an octave that would contain most of the spectra, accounting for the longer period of the *lin-28(0)* strain.

The analytic signal of each preprocessed gene was then computed using the Hilbert Transform. The instantaneous phase can be computed as the argument of the analytic signal. The instantaneous frequency was computed by differentiating numerically the unwrapped instantaneous phase (functions *angle, unwrap* and *diff* from the Numpy library, version 2.3.1). The instantaneous period is the reciprocal of the instantaneous frequency. The distribution of instantaneous periods was computed for each timepoint for the whole set of oscillating genes in each condition.

#### Small RNA-seq

RNA integrity was validated using RNA Nano Chip with Agilent 2100 Bioanalyzer and sequencing libraries were generated using CleanTag Small RNA Library Preparation Kit (Trilink) according to the manufacturer’s instructions. The library was sequenced on Illumina HiSeq2500, as 50 cycles single-end reads. The small RNA-seq fastq files were adapter trimmed (TGGAATTCTCGGGTGCCAAGG) at the 3′ end using the function preprocessReads() from the QuasR package. Sequencing reads were mapped to the *C. elegans* genome (ce10) using the qAlign() function from the QuasR package. For miRNA quantification, gene annotations from miRBase v22 were used. We extended the genomic coordinates by ±3 bp (upstream and downstream) of the 5′ end of each miRNA and counted all the reads that started within those regions. Read counts were normalized to the mean mapped library size and the normalized values were then log_2_-transformed with a pseudocount of 8. The data obtained from this timecourse (TC2) were used to generate the expression profiles of different miRNAs in Fig. 1F and S2.

To determine differentially expressed miRNAs in *lin-28(0)* animals (Fig. 1G), we used a published small RNA seq dataset (Nahar *et al*., 2024). In this dataset, a part (0-23 h) of the long developmental timecourse TC2 was resequenced for deeper coverage and the sequencing reads were combined with the reads from the whole timecourse sequencing and further processed as described in Nahar et al., 2024. Differentially expressed miRNAs were identified using the R package edgeR, treating time points 3-6 h as pseudoreplicates, with batch correction applied for time points. miRNAs with low expression were filtered out using filterByExpr. Quasi-likelihood F-tests were performed to assess differential expression, and miRNAs with a false discovery rate (FDR) < 0.01 were considered significantly differentially expressed.

### LIN28 RNA-immunoprecipitation followed by sequencing (RIP-seq)

#### Sample collection and lysis

The experiment was performed with non-transgenic wild-type animals (N2; control) and *lin-28::large_tag* animals (HW2805). Approximately 4.5 million of synchronized L1s of each genotype were plated on peptone-enriched 2% NGM agar XL plates seeded with *E. coli* OP50 bacteria (∼ 500,000 L1s per plate) and grown for 6 hours at 25°C. Animals were harvested, washed three times with 10 ml M9 buffer, washed once with 8 ml of 0.1 M KCl, resuspended in ∼1.5 ml of 0.1 M KCl and transferred to a 1.5-ml Eppendorf tube. Samples were spun down, the remaining liquid was aspirated and dry pellets were snap-frozen in liquid nitrogen. Upon thawing, animal pellets were lysed in an approximately equal volume of extraction buffer (50 mM HEPES/KOH (pH 7.4 at 4°C), 150 mM KCl, 5 mM MgCl_2_, 0.1% (v/v) Triton X-100, 10% (w/v) glycerol, 1 mM PMSF, 1 tablet/10 ml cOmplete Mini Protease Inhibitor Tablets (EDTA-free; Roche, 11836170001), 200 U/ml RNaseOUT Recombinant Ribonuclease Inhibitor (Thermo Fisher Scientific, 10777019)), with mortar and pestle in the presence of liquid nitrogen. Lysates were cleared by centrifugation at 12,000g for 20 min at 4°C. The supernatant was transferred to a fresh Eppendorf tube and kept on ice.

#### RNA-Immunoprecipitation (RIP)

Total protein concentration was measured using Pierce BCA Protein Assay Kit (Thermo Fisher Scientific, Cat.no. 23225 and 23227). Anti-GFP IPs were performed by incubating 3 mg total protein with 50 μl of Dynabeads Protein G (Thermo Fisher Scientific, 10003D) bound to anti-GFP antibody (5 μg/IP; Abcam, ab290) in a total volume of 1 ml for 3 hours at 4°C on a rotating wheel. Beads were washed five times for five minutes in 1 ml of extraction buffer without protease and RNase inhibitors, before extracting the bound RNA by directly adding TRI Reagent to the beads and snap-freezing in liquid nitrogen. For each condition, three IPs were performed in parallel as technical replicates. Three aliquots of input lysate each corresponding to 100 μg total protein were snap-frozen in liquid nitrogen upon addition of TRI Reagent for extraction of the input RNA.

#### RNA extraction and sequencing

Upon thawing, samples were incubated at room temperature for 10 min shaking at 300 rpm before transferring the supernatant without the beads to a new Eppendorf tube. RNA was extracted using Direct-zol RNA Microprep kit (Zymo Research; R2062) according to the manufacturer’s protocol including the on-column DNase digestion. RNA integrity was validated using RNA Nano Chip with Agilent 2100 Bioanalyzer and sequencing libraries were generated using Illumina Stranded mRNA-seq protocol. The library was sequenced using NovaSP, 50 cycles pair-end reads.

#### RIP-seq analysis

Sequencing reads were mapped to the *C. elegans* reference genome (ce11) with the R package QuasR using the spliced alignment algorithm HISAT2 (Gaidatzis *et al*., 2015). Protein-coding exons were defined based on the exon annotation file from WormBase (WS270) and read mappings to the corresponding genes were counted using qCount function from the QuasR package.

To determine differentially enriched mRNAs in LIN28 IP relative to untagged control (N2) IP (Fig. 1B), we used the R package edgeR. Genes with low expression were filtered out using filterByExpr. Quasi-likelihood F-tests were performed to assess differential enrichment, and genes with a false discovery rate (FDR) < 0.01 were considered significantly differentially enriched.

To examine whether *pri-let-7* is enriched in LIN28 IP relative to untagged control (N2) IP (Fig. 1E), *pri-let-7* locus was defined with genome coordinates using the Grange function (GRanges(seqnames = Rle(“chrX”),ranges = IRanges(14743780, end = 14744173),strand=“-”)), starting right downstream of the *pre-let-7* stem-loop up until the *C05G5.7* coding sequence. Reads mappings to this region were counted using qCount function from the QuasR package and the counts were normalized to the mean mapped library size. The normalized values were then log_2_-transformed with a pseudocount of 1. The statistical significance of differences in pri-let-7 enrichment between the averages of the two input samples and the averages between the two IP samples was assessed using an unpaired two-sided *t*-test.

#### Luciferase assay

To assess the molting cycle progression of individual animals, we employed a luciferase assay as described previously (Meeuse *et al*., 2020; Olmedo *et al*., 2015). Gravid adults were bleached and single embryos expressing luciferase from a constitutive and ubiquitous promoter (transgene *xeSi312*) were transferred by pipetting to a white, flat-bottom, 384-well plate (Berthold Technologies, 32505). Embryos were allowed to develop in 90 μl S-Basal medium containing *E. coli* OP50 bacteria (OD600 = 1) and 100 μM Firefly D-Luciferin (p.j.k., 102111). Plates were sealed with Breathe Easier sealing membrane (Diversified Biotech, BERM-2000). Luminescence was measured every 10 min for 0.5 seconds for a total duration of 52-96 h (depending on the purpose of the experiment) using a luminometer (Berthold Technologies, Centro XS3 LB 960). The luminometer was placed in a temperature-controlled incubator set to 18.5°C, which translated to an effective temperature of 22.5°C experienced by the animals inside the luminometer.

Luminescence intensity traces of individual animals were further analyzed using a Python-based custom code developed by L.J.M.M.. Hatch, as well as both molt entries and exits, were automatically annotated using a small convolutional neural network and further refined manually. Molts were defined as periods of stable low signal, preceded and followed by a sudden drop and rise in luminescence intensity, respectively. Significant differences in total durations of larval development (Fig. 7A) were determined by an unpaired two-sided *t*-test. *P*-values of all pair-wise comparisons of L3 durations were determined by a non-parametric Dunn’s test with Holm correction for multiple testing (Fig. 7D, 7E, 7F, S7).

#### Examination of animals carrying lethal alleles balanced with *xeEx365* array

Several strains examined in this work (Table S1) carry the lethal *let-7(xe150)* allele, which causes vulva bursting shortly upon the exit from the fourth molt. To maintain these strains, they were balanced with the extrachromosomal *xeEx365* array that contains a *let-7* transgene and several mCherry reporters (*xeEx365[let-7p::let-7::SL1_operon_GFP, unc-119 (+); Prab-3::mCherry; Pmyo-2::mCherry; Pmyo-3::mCherry]*). This array segregates randomly during meiosis, resulting in incomplete inheritance, with only a subset of progeny carrying the array and thus surviving into adulthood. In order to examine the molting cycle of animals with lethal genotypes, animals were retrieved from the luciferase assay plate after the assay was finished. They were visually inspected for the presence of the mCherry-marked balancer array using a fluorescent binocular microscope and only the luminescence intensity traces of animals not carrying the balancer array were included in the analysis.

#### Examination of the 4M* phenotype in *lin-28(0); lin-46(0)* animals

To examine the partially penetrant and liquid culture-specific aberrant fourth molt phenotype of *lin-28(0); lin-46(0)* animals (HW3504), synchronized L1s were loaded to the luciferase assay plate. The reason for using synchronized L1s instead of embryos was to achieve a synchronously developing population to enrich for animals at the desired developmental stage at the time of examination. The luciferase assay was run for 52 hours. Afterwards, the luminescence intensity traces of individual animals were examined and classified into two groups – animals that underwent a normal fourth molt and animals that exhibited the 4M* phenotype (a steep increase in luminescence intensity instead of the typical signal drop at the time of the expected fourth molt). Fifteen animals from each group were retrieved from the plate, mounted onto two separate 2% (w/v) agarose pads and immobilized in 10 mM levamisole. Differential Interference Contrast (DIC) images were acquired with a Zeiss Axio Observer Z1 microscope using the Zen 2 software. Selections of regions and image processing were performed with Fiji (Schindelin *et al*, 2012).

#### Confocal imaging

To assess the time of LIN-46 protein accumulation, animals were grown on 2% NGM agar OP50 plates at 25°C for the indicated time before mounted on a 2% (w/v) agarose pad with a drop of 10 mM levamisole solution. Animals were imaged on an upright microscope Axio Imager M2 (Zeiss) equipped with CSU W1 with Dual T2 spinning disk confocal scanning unit (Yokogawa) driven by Visiview 6.0 (Visitron). Differential Interference Contrast (DIC) and fluorescent images were acquired with a 40x/1.3 oil immersion objective (EC Plan Neofluar). Using the Fiji software, images were processed after selecting representative regions.

#### Western blotting

Synchronized L1s were plated on 10 cm 2% NGM agar OP50 plates, grown at 25°C and harvested at 3 and 6 h (20,000 animals per time point) in M9 buffer, washed twice with M9 buffer, resuspended in ∼1 ml of 0.01% Tween-20 (Sigma-Aldrich, P9416) in M9 buffer and transferred to 1.5 ml Eppendorf tubes. Samples were spun down and dry pellets were snap-frozen in liquid nitrogen. Lysates were prepared by boiling (5 min, 95°C) and sonication (Bioruptor Plus, 13 cycles, 30 s on/60 s off at 4°C) in SDS lysis buffer (63 mM Tris-HCl (pH 6.8), 5 mM DTT, 2% SDS, 5% sucrose) and cleared by centrifugation. Proteins were separated by SDS-PAGE on a 4-12% Bis-Tris gel (Thermo Fisher Scientific, NP0335BOX) in MOPS running buffer (Thermo Fisher Scientific, NP0001). Samples were run at 80 V for ∼30 min and then at 150 V until the desired protein separation was achieved. 40 μg of protein extract was loaded per well and PageRuler Prestained Protein Ladder (10 to 180 kDa; Thermo Fisher Scientific, 26616) was used as a molecular weight reference. Proteins were transferred to a PVDF membrane (Bio-Rad, 1620177) by semi-dry blotting at 20 V for 1 hour (transfer buffer: 250 mM Tris, 1.9 M glycine, 0.5% SDS + 20% methanol). The membrane was blocked with 5% (w/v) milk in TBST (0.1% Tween-20 in TBS) for 1 h shaking (150 rpm). LIN28 was detected using polyclonal rabbit anti-LIN28 antibody (custom made by GenScript, R13880; raised against full-length recombinant His6::TEV::LIN28a; 1:1,000), followed by polyclonal anti-rabbit HRP-conjugated antibody (Santa Cruz Biotechnology, sc-2004; 1:5,000). Antibodies were diluted in 0.5% (w/v) milk in TBST and incubated with the membrane for 1 h shaking at 150 rpm at room temperature. ECL Detection Reagent (Cytiva, RPN2232) and Amersham Imager 680 were used for detection. After LIN28 detection, the membrane was stripped (Restore PLUS Western Blot Stripping Buffer; Thermo Fisher Scientific, 46430), washed with TBST, blocked as described above and incubated with monoclonal mouse anti-Actin, clone C4 (MilliporeSigma, MAB1501; 1:7,500) and anti-mouse HRP-conjugated antibody (Cytiva, NXA931; 1:5,000) in 0.5% (w/v) milk in TBST for 1 h shaking at 150 rpm at room temperature, followed by Actin detection as described above.

#### Examination of the protruding vulva and precocious alae phenotypes upon molt exit

To assess the vulva morphology and the presence of alae, synchronized L1s of indicated genotypes were grown on 2% NGM agar OP50 plates at either 20°C or 25°C and inspected with a binocular microscope. When ∼50% of the animals entered the respective molt (Table S3), as indicated by the lack of movement and pharynx pumping, ∼35 animals were individually transferred to small plates and inspected at regular intervals (every ∼20 min) for the signs of molt completion, as indicated by resumed movement, pharynx pumping and the presence of the old shed cuticle on the plate. The first ten animals that exited the molt were mounted onto a 2% (w/v) agarose pad, immobilized in 10 mM levamisole and the phenotypes were examined. Afterwards, ten more animals that exited the molt in the meantime were processed in the same way. Differential Interference Contrast (DIC) images were acquired with a Zeiss Axio Observer Z1 microscope using the Zen 2 software. Selections of regions and image processing were performed with Fiji.

#### Quantification of the protruding vulva phenotype

Synchronized L1s were plated on 2% NGM agar OP50 plates and grown at 20°C for 52 h, before animals exhibiting either normal or protruding vulva were counted using a binocular microscope. Four plates per condition were examined (N = 4), with > 230 animals scored per condition. Significant differences to wild type were determined by a Dunnett’s test.

#### Quantification of the vulva bursting phenotype

Synchronized L1s were plated on 2% NGM agar OP50 plates and grown at 25°C for 48 h, before animals exhibiting either normal or burst vulva were counted using a binocular microscope. Three plates per condition were examined (N = 3), with 240 animals scored per condition (n = 240).

## Acknowledgements

We thank Benjamin Towbin, Rajani Gudipati and Julien Orgül for generating strains used in this study, Iskra Katic for microinjections, Sandra Mühlhäusser for producing recombinant LIN28 for antibody production, Victor Ambros for providing strains, Alex Smith for statistical analysis support and Chiara Azzi for experimental support. We thank Marc Bühler and Iskra Katic for their comments on the manuscript.

## Author Contributions

J.B.: Conceived project, designed, executed and analyzed experiments. Wrote manuscript.

A.W.: Designed and executed experiments, analyzed data.

D.G.: Analyzed data.

L.J.M.M.: Analyzed data.

H.G.: Conceived and supervised project, analyzed data. Wrote manuscript. Acquired funding.

## Funding

The research leading to these results has received funding from the European Research Council (ERC) under the European Union’s Horizon 2020 research and innovation program (Grant agreement No. 741269), the Swiss National Science Foundation (SNF, #310030_188487 & 310030_219271), and the Novartis Research Foundation (through the FMI) (to H.G.).

## Conflict of Interes

The authors declare that they have no conflict of interest.

## Data Accessibility

All RNA-seq data are accessible at NCBI’s Gene Expression Omnibus (Edgar *et al*, 2002) under SuperSeries accession number GSE315957 at https://www.ncbi.nlm.nih.gov/geo/query/acc.cgi?acc=GSE315957

